# Resident Macrophages are Locally Programmed for Silent Clearance of Apoptotic Cells

**DOI:** 10.1101/155010

**Authors:** Allison W. Roberts, Bettina L. Lee, Jacques Deguine, Shinu John, Mark J. Shlomchik, Gregory M. Barton

## Abstract

Although apoptotic cells (ACs) contain nucleic acids that can be recognized by Toll-like receptors (TLRs), engulfment of ACs does not initiate inflammation in healthy organisms. To better understand this phenomenon, we identified and characterized macrophage populations that continually engulf ACs in several distinct tissues. These macrophages share characteristics compatible with immunologically silent clearance of ACs, including high expression of AC recognition receptors, low expression of TLR9, and reduced TLR responsiveness to nucleic acids. When removed from tissues these macrophages lose many of these characteristics and generate inflammatory responses to AC-derived nucleic acids, suggesting that cues from the tissue microenvironment are required to program macrophages for silent AC clearance. We show that KLF2 and KLF4 control expression of many genes within this AC clearance program. Coordinated expression of AC receptors with genes that limit responses to nucleic acids may represent a central feature of tissue macrophages that ensures maintenance of homeostasis.

## INTRODUCTION

Induction of inflammatory responses upon recognition of nucleic acids is a critical feature of the innate immune system. Nucleic acids are recognized by multiple pattern recognition receptors (PRRs), including endosomal Toll-like receptors (TLRs) specific for DNA (TLR9) and various forms of RNA (TLR3, TLR7, TLR8, and TLR13)(Barbalat et al., 2011). This strategy affords broad recognition of multiple pathogen classes, and its failure can render the host susceptible to infection by a variety of pathogens (Barrat et al., 2016). However, the cost of this broad recognition is the potential for inappropriate responses to self-derived nucleic acids, which can lead to autoimmunity or autoinflammatory diseases (Sharma et al., 2015).

Multiple mechanisms function to limit the likelihood of responses against self by TLRs that recognize nucleic acids. For instance, TLR9 preferentially recognizes DNA that contains unmethylated CpG dinucleotides, and these motifs are more frequent in microbial DNA than mammalian DNA (Coch et al., 2009; Krieg et al., 1995; Yasuda et al., 2009). In addition, TLR9 and TLR7 are localized to endosomes which limits access to extracellular self-derived DNA and RNA while allowing responses to pathogens that access the endosome (Barton et al., 2006; Mouchess et al., 2011). Mechanisms that bypass this compartmentalization can disrupt homeostasis. For example, the generation of immune complexes containing nucleic acids can lead to Fc receptor-mediated uptake of endogenous nucleic acids, activation of endosomal TLRs, and subsequent autoimmune responses (Boulé et al., 2004; Leadbetter et al., 2002; Means et al., 2005).

Avoidance of self nucleic acid recognition during clearance of apoptotic cells (ACs) presents additional challenges. First, the volume of cargo that must be cleared is immense; it has been estimated that millions of cells die by apoptosis in the human body every day (Fond and Ravichandran, 2016). If clearance is disrupted, accumulation of ACs can lead to immune stimulation and, eventually, autoimmune disease (Asano et al., 2004; Baumann et al., 2002; Hanayama et al., 2004). Second, professional phagocytes, such as macrophages and dendritic cells, that engulf ACs express TLRs capable of nucleic acid recognition. Third, after recognition by a variety of phagocytic receptors (Miyanishi et al., 2007; Park et al., 2008; Scott et al., 2001), ACs traffic to phagosomes, the same organelles that house nucleic-acid sensing TLRs. Thus, the compartmentalization of TLR9 and TLR7 is not sufficient to explain the lack of response to the nucleic acids within ACs. Nevertheless, AC-derived nucleic acids do not typically initiate inflammatory responses. This avoidance is generally attributed to AC-induced expression of anti-inflammatory mediators. Largely through *in vitro* studies, it has been shown that ACs can induce anti-inflammatory cytokine production as well as cell autonomous anti-inflammatory signaling pathways in phagocytes (A-Gonzalez et al., 2009; Freire-de-Lima et al., 2006; McDonald et al., 1999; Rothlin et al., 2007; Tassiulas et al., 2007). However, *in vivo* AC clearance is a constant process, and it remains unclear how the innate immune system balances induction of anti-inflammatory responses while maintaining the ability to respond to pathogens.

It has been proposed that tissue-resident macrophages are important mediators of AC clearance (Fond and Ravichandran, 2016). Several macrophage populations have been demonstrated to engulf ACs injected into mice (McGaha et al., 2011; Miyake et al., 2007; Uderhardt et al., 2012; Wang et al., 2008). In addition, it has recently been demonstrated that apoptotic intestinal epithelial cells are engulfed by a DC subset and two macrophage populations in the intestine (Cummings et al., 2016). However, the identities of the cells that clear ACs from most tissues in the steady state remain unclear. This issue is particularly interesting in light of recent evidence that macrophages from different tissues are quite heterogeneous (Gautier et al., 2012b). Although macrophage precursors express a core macrophage transcriptional program, macrophages diversify after colonizing tissues (Mass et al., 2016). Much of this diversity is controlled by local signals from tissues that induce gene expression and dictate the phenotype and function of resident macrophages (Gosselin et al., 2014; Lavin et al., 2014; Okabe and Medzhitov, 2014). It remains unclear whether this environmental programming induces heterogeneity in the ability of different resident macrophage populations to clear ACs or influences responses to ACs.

To investigate how self-tolerance is maintained during AC-clearance we have identified macrophage populations in several tissues that efficiently engulf ACs *in vivo* at steady state. These macrophages are programmed by their tissue environment to limit responses to low doses of nucleic acid TLR ligands and to lack expression of TLR9. We identify two transcription factors, KLF2 and KLF4, as critical regulators of this AC-clearance program. KLF2 and KLF4 induce expression of genes that facilitate AC uptake as well as negatively regulate TLR signaling. Coordinated expression of receptors that promote AC clearance with genes that limit innate responses to the nucleic acids within ACs may represent a central feature of tissue macrophages that ensures maintenance of homeostasis in the face of constant AC clearance.

## RESULTS

### Identification of macrophage populations that clear apoptotic cells *in vivo*

To investigate how tolerance to AC-derived ligands is maintained we first sought to identify cells that clear ACs under homeostatic conditions *in vivo*. We generated mixed bone marrow chimeric mice: C57BL/6 (WT) bone marrow expressing the congenic marker CD45.2 was mixed with bone marrow expressing the congenic marker CD45.1 and the fluorescent protein tdTomato and injected into irradiated WT recipients (Figure 1A). In these chimeric mice the only cells that genetically encode for tdTomato also express CD45.1. Therefore, we reasoned that these mice could be used to identify cells involved in dead cell clearance, as tdTomato^+^CD45.2^+^CD45.1^−^ cells must have acquired tdTomato by engulfing a CD45.1^+^ cell. The use of congenic markers allowed us to exclude any tdTomato^+^CD45.2^+^CD45.1^+^ cells resulting from cell-cell fusion. A recent study used a similar system to track engulfment of GFP+ intestinal epithelial cells and observed a low level of constitutive engulfment, which they also attributed to steady-state clearance of ACs (Cummings et al., 2016).

**Figure 1.**
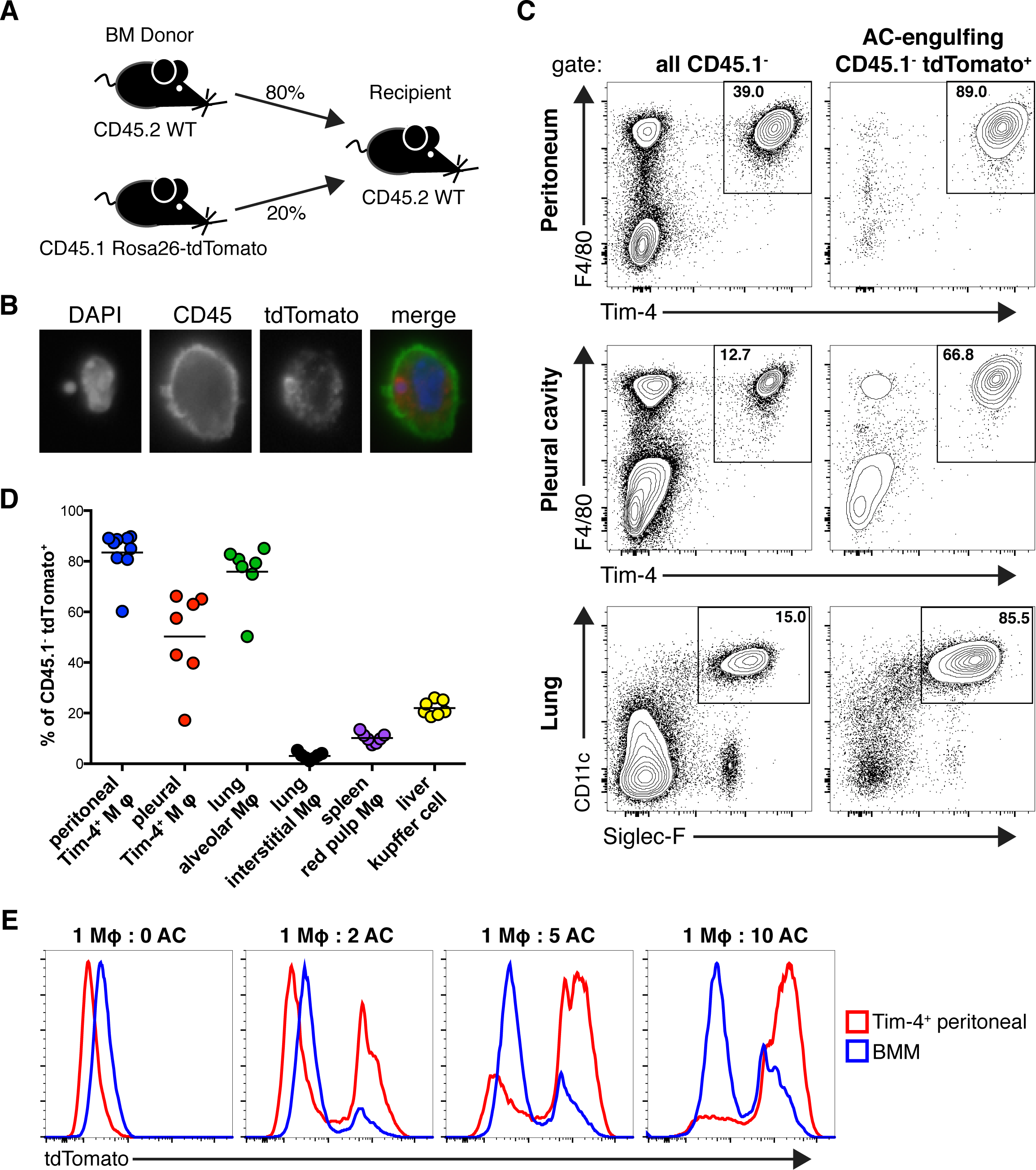
Identification of tissue resident macrophage populations that clear apoptotic cells *in vivo*. **(A)** Diagram of bone marrow chimeras used to identify AC-engulfing cells at steady state. Congenically marked CD45.1^+^bone marrow cells expressing tdTomato were mixed with wild type CD45.2^+^ bone marrow cells and injected into an irradiated wild type recipient. Tissues were analyzed after 10 weeks of reconstitution. **(B)** Detection of cells that contain tdTomato^+^ cell corpses. Representative immunofluorescence picture of AC-engulfing cell isolated from chimeras depicted in (A). Cells were stained with DAPI and antibody against CD45. **(C)** Tim-4^+^ pMacs, alveolar macrophages, and Tim-4^+^ pleural cavity macrophages are adept at engulfing ACs *in vivo.* Representative flow cytometric analyses of tissues from bone marrow chimeras. Cells were gated on CD45.1^−^ cells or AC-engulfing cells (CD45.1^−^ tdTomato^+^), as indicated. Data are representative of at least three independent experiments. **(D)** Quantification of AC engulfment by the indicated macrophage populations in tissues. Data are the combination of three independent experiments with total n = 7-9 for all groups. **(E)** Tim-4^+^ pMacs are adept at engulfing ACs. CD45.2^+^ Tim-4^+^ pMacs and CD45.2^+^ BMMs were incubated with CD45.1 + tdTomato^+^ ACs at indicated ratios and analyzed by flow cytometry. Data are representative of three independent experiments. See also Figure S1.

After ten weeks of reconstitution, tissues from these mice were harvested and analyzed by flow cytometry. After gating out CD45.1^+^ cells that express TdTomato, we identified a population of tdTomato^+^ cells that was not present in control chimeras generated with CD45.1^+^ bone marrow lacking tdTomato (Figure S1A). Since the CD45.1^−^ cells do not genetically encode for tdTomato themselves, we reasoned that they had engulfed tdTomato-expressing ACs. Of course, it is formally possible that they acquired tdTomato by engulfing necrotic cells or cell debris, but one would not expect significant necrotic death under homeostatic conditions. Immunofluorescence microscopy of cells from the tissues of chimeric mice confirmed the presence of cells containing tdTomato^+^ cell corpses, as shown by the colocalization of tomato and DNA within WT cells (Figure 1B).

After analyzing multiple tissues, we identified three tissue-resident macrophage populations as the primary engulfers of dead hematopoietic cells at steady state in their respective tissues: peritoneal macrophages (pMacs) that express the AC recognition receptor Tim-4, pleural cavity macrophages that express Tim-4, and lung alveolar macrophages, which do not express Tim-4 (Figures 1C, 1D, and S1B). To confirm the results of the bone marrow chimera system we independently established that these populations efficiently engulf labeled ACs *in vivo* and *in vitro* (Figures 1E, S1C, and S1D). In other tissues analyzed, we did not identify a single macrophage population that was the primary engulfer of dead hematopoietic cells. However, we did identify several macrophage populations that engulf dead cells to a lesser extent than the three populations described above: CD11b^+^ interstitial macrophages in the lung, red pulp macrophages in the spleen, and Kupffer cells in the liver (Figures 1D, S1B and S1E). We did not identify a macrophage population that engulfed dead hematopoietic cells in skin draining lymph nodes, but noted that CD11c^mid^ migratory dendritic cells represented about 25% of CD45.1^−^tdTomato^+^cells, suggesting that they may acquire dead cells in the skin and then migrate to the lymph nodes (Figures S1B and S1E). In several organs, including the spleen and pleural cavity, we observed a B220^+^ population that engulfed dead hematopoietic cells. These cells are likely B lymphocytes that have engulfed apoptotic bodies as has previously been described for B-1 cells (Rodriguez-Manzanet et al., 2010). We also identified a dead cell engulfing CD45^−^ population in the liver, which is likely either hepatocytes or endothelial cells that have previously been described to engulf ACs (Dini et al., 2002). The differences in CD45.1^−^tdTomato^+^cells between organs were not due to large differences in the number of available ACs, as the percentage of activated Caspase-3/7^+^tdTomato^+^cells was similar in all organs we examined, except the liver which had more ACs (Figure S1F).

### AC-engulfing macrophage populations do not express TLR9

Having measured which cell populations engulf dead cells under homeostatic conditions *in vivo*, we chose to further characterize the peritoneal, pleural, and alveolar macrophages because these populations represented the clearest examples of cells dedicated to AC clearance in our system. We also noted that these three macrophage populations are localized to body cavities and wondered whether they share features that enable silent clearance of dead cells at these sites. As a first step we analyzed expression of TLR9 and TLR7 using two lines of knock-in reporter mice, TLR9KI and TLR7KI (Figure 2A). Examining endogenous TLR expression is notoriously challenging due to a lack of reliable antibodies. These new reporter mice enable the analysis of endogenous TLR expression levels with C-terminal epitope tags and fluorescent protein driven by internal ribosome entry sequences downstream of the TLR coding sequence. Intriguingly, the three AC-engulfing macrophage populations described above did not express TLR9 (Figure 2B). We confirmed the lack of TLR9 protein by immunoblot (Figure 2C). Other macrophage populations, such as red pulp macrophages, Kupffer cells, lung CD11b^+^ interstitial macrophages, and F4/80^mid^ peritoneal and pleural macrophages did express TLR9 (Figures 2B and S2A). F4/80^hi^Tim-4^−^ peritoneal and pleural macrophages did not express TLR9, but it remains unclear whether these macrophages represent truly independent populations from the Tim-4^+^ populations in these sites. TLR7 was expressed in most macrophages studied except for a proportion of the F4/80^mid^ peritoneal and pleural macrophage populations.

**Figure 2.**
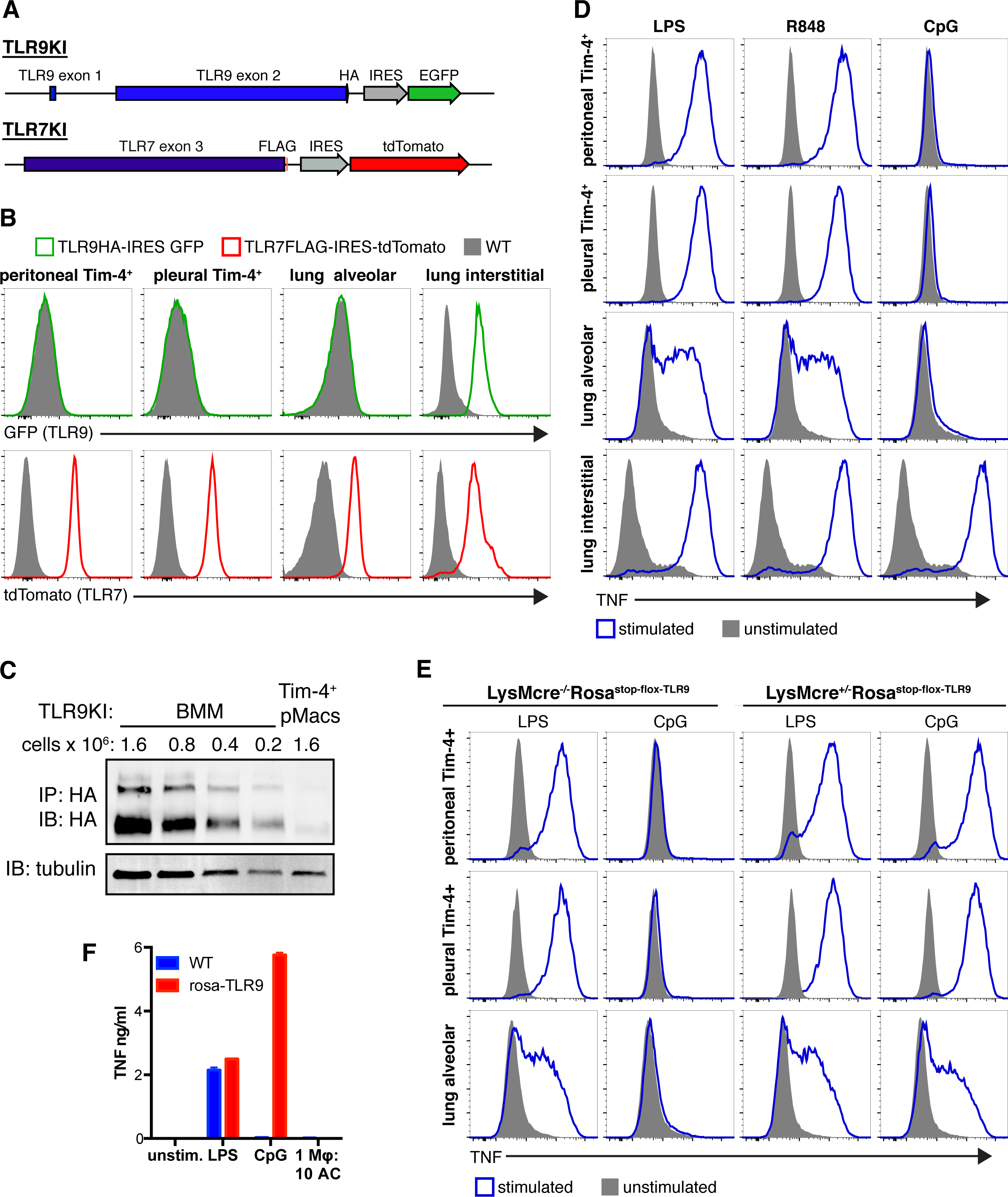
AC-engulfing macrophages do not express TLR9. **(A)** Diagram of reporter mice generated to examine TLR expression. **(B)** AC-engulfing macrophages do not express TLR9. Representative histograms of fluorescent protein expression by gated macrophages harvested from TLR9KI, TLR7KI, or WT mice. Peritoneal and pleural Tim-4^+^ macrophages are gated as live F4/80^+^Tim-4^+^ cells. Alveolar macrophages are gated as live SiglecF^+^CD11c ^+^CD64 +CD11b ^−^ cells. Lung interstitial macrophages are gated as live CD11b^+^F4/80^+^MHCII^+^ cells. Data are representative of at least three independent experiments with n = 2 per experiment. **(C)** Tim-4^+^ pMacs and BMMs from TLR9KI mice were analyzed for TLR9 protein expression. Cell lysates were immunoprecipitated with anti-HA resin and eluted protein was analyzed by anti-HA western. The two bands represent ER localized TLR9 (larger band) and endosomal cleaved TLR9 (smaller band). An anti-tubulin immunoblot was performed on the same lysates. Relative densities normalized to tubulin are 1 for 0.2e6 BMM and 0.11 for 1.6e6 Tim-4^+^ pMacs. Data are representative of at least three independent experiments. **(D)** AC-engulfing macrophages do not respond to TLR9 ligands. Macrophages from WT mice were stimulated *ex vivo* with TLR ligands. Macrophages were gated as in (A). TNF production was measured by ICS. Data are representative of at least three independent experiments with n = 2 per experiment. **(E)** When forced to express TLR9 AC-engulfing macrophages respond to CpG ODN. Macrophages from LysMcre^+/-^Rosa^stop-flox-TLR9^ or littermate control LysMcre^−/-^Rosa^stop-flox-TLR9^ mice were stimulated *ex vivo* with TLR ligands. TNF production was measured by ICS. Data are representative of at least three independent experiments. **(F)** Tim-4^+^ pMacs forced to express TLR9 do not generate responses to ACs. Isolated rosa-TLR9 Tim-4^+^ pMacs were stimulated with TLR ligands and ACs. TNF was measured by CBA. Data are representative of at least three independent experiments. See also Figure S2.

We confirmed that the three AC-engulfing macrophage populations lack TLR9 by stimulating macrophages *ex vivo* with TLR ligands. In agreement with the results from the reporter mice, these macrophages produced TNF in response to stimulation with LPS (TLR4 ligand) or R848 (TLR7 ligand) but not CpG oligonucleotides (ODN), a synthetic TLR9 ligand (Figures 2D and S2B). CD11b^+^ interstitial macrophages and red pulp macrophages generated robust responses to TLR9 stimulation. Interestingly, while Kupffer cells in the liver did express TLR9 and generated responses to TLR9 stimulation, these responses were muted compared to responses to other TLR ligands (Figure S2B). Kupffer cells also represented a larger percent of the AC-engulfing cells in the liver compared to red pulp macrophages or lung interstitial macrophages (Figure 1D). Therefore, Kupffer cells are intermediate in both engulfment of ACs and responses to TLR9 stimulation, suggesting that there may be an inverse correlation between AC engulfment ability and expression of TLR9. Altogether, these data led us to hypothesize that tight regulation of TLR9 expression is one mechanism used to avoid responses to self-derived DNA.

### Forcing expression of TLR9 in AC-engulfing macrophages only slightly enhances inflammation during pristane-induced lupus

To investigate whether lack of TLR9 expression in Tim-4+ pMacs, pleural cavity macrophages, and alveolar macrophages is needed to maintain self-tolerance we forced these macrophages to express TLR9. We generated mice that ectopically express TLR9 in all cells (rosa-TLR9) or in LysM^+^ cells (LysMcre+/-Rosa^stop-flox-TLR9^) by crossing mice in which TLR9 is inserted behind a floxed stop cassette in the Rosa26 locus to EIIA-cre or LysM-cre mice, respectively. Tim4^+^ pMacs, pleural cavity macrophages, and alveolar macrophages from LysMcre^+/-^Rosa^stop-flox-TLR9^ mice responded to CpG ODN (Figure 2E). This result confirms that AC-engulfing macrophages control responses to TLR9 ligands through TLR9 expression and rules out additional mechanisms such as altered receptor trafficking or repression of cytokine production downstream of signaling. However, we were surprised to find that despite this forced TLR9 responsiveness, Tim-4^+^ pMacs from rosa-TLR9 mice did not respond to ACs *in vitro* (Figure 2F). This lack of response was not due to reduced AC engulfment (Figure S2C).

When housed unperturbed in our specific pathogen-free mouse facility, LysMcre^+/-^Rosa^stop-flox-TLR9^ mice showed no evidence of inflammatory disease over twelve months (data not shown), demonstrating that forced expression of TLR9 in AC-engulfing macrophages is not sufficient to disrupt homeostasis. We next considered whether restricted expression of TLR9 might be critical under conditions where there is increased cell death *in vivo*. To this end we employed the pristane-induced model of lupus, in which intraperitoneal injection of pristane oil induces substantial apoptosis in the peritoneal cavity followed by recruitment of inflammatory cells and development of a lupus-like disease, including production of anti-nuclear antibodies ((Calvani et al., 2005)). It has previously been demonstrated that TLR7 and TLR9 play roles in the pathogenesis of this disease model (Lee et al., 2008; Summers et al., 2010). We compared LysMcre^+/-^Rosa^stop-flox-TLR9^ and LysMcre^−/-^Rosa^stop-flox-TLR9^ littermates at two weeks and nine months post injection. LysMcre^+/-^Rosa^stop-flox-TLR9^ mice had small increases in the number of cells recruited into the peritoneum at two weeks, including neutrophils and Ly6C^+^ monocytes (Figures S2D and S2E). After nine months LysMcre^+/-^ Rosa^stop-flox-TLR9^ mice still demonstrated a small increase in peritoneal cell numbers, including CD11b^+^ cells, T cells, and neutrophils (Figures S2D and S2F). Sera from these mice also contained increased anti-dsDNA IgG (Figure S2G).

Altogether, the overexpression of TLR9 in LysM^+^cells slightly enhanced inflammation in the pristane model of lupus; however, the subtlety of these differences clearly indicated that additional mechanisms must be limiting tissue macrophage responses to nucleic acids within ACs.

### Tissue environments program resident macrophages for silent clearance of ACs

Based on the modest effect of TLR9 overexpression in tissue macrophages *in vivo*, and the fact that TLR9 overexpression is not sufficient to induce TNF production in response to ACs *in vitro*, we reasoned that these cells must employ additional mechanisms to prevent responses to ACs. Many recent studies have demonstrated that tissue-resident macrophages are programmed by cues from their environment to express specific transcription factors that shape their identity and function (Gautier et al., 2012b; Gosselin et al., 2014; Lavin et al., 2014; Okabe and Medzhitov, 2014). We hypothesized that AC-engulfing macrophages may be subject to similar programming of which inhibition of TLR9 expression is only one facet. To investigate this possibility, we compared isolated Tim-4^+^ pMacs that had been cultured *ex vivo* overnight to macrophages that had been cultured *ex vivo* for three days. Remarkably, the macrophages cultured for three days not only gained responsiveness to CpG ODN but also began responding to ACs (Figures 3A and S3A). Similar results were obtained with alveolar macrophages and Tim-4^+^ pleural macrophages (Figures 3B and 3C). This altered response corresponded with increased expression of TLR9 protein (Figure 3D) and was not due to changes in AC uptake (Figure S3B). Tim-4^+^ pMacs from mice deficient for TLR signaling adaptors MyD88 and TRIF or for UNC93B1, a trafficking chaperone required for endosomal TLR function, failed to respond to ACs, implicating endosomal TLRs in the response to ACs (Figures 3E and S3B). In contrast, bone marrow-derived macrophages (BMMs) failed to eat ACs efficiently and did not produce TNF (Figures S3C and 1E).

**Figure 3.**
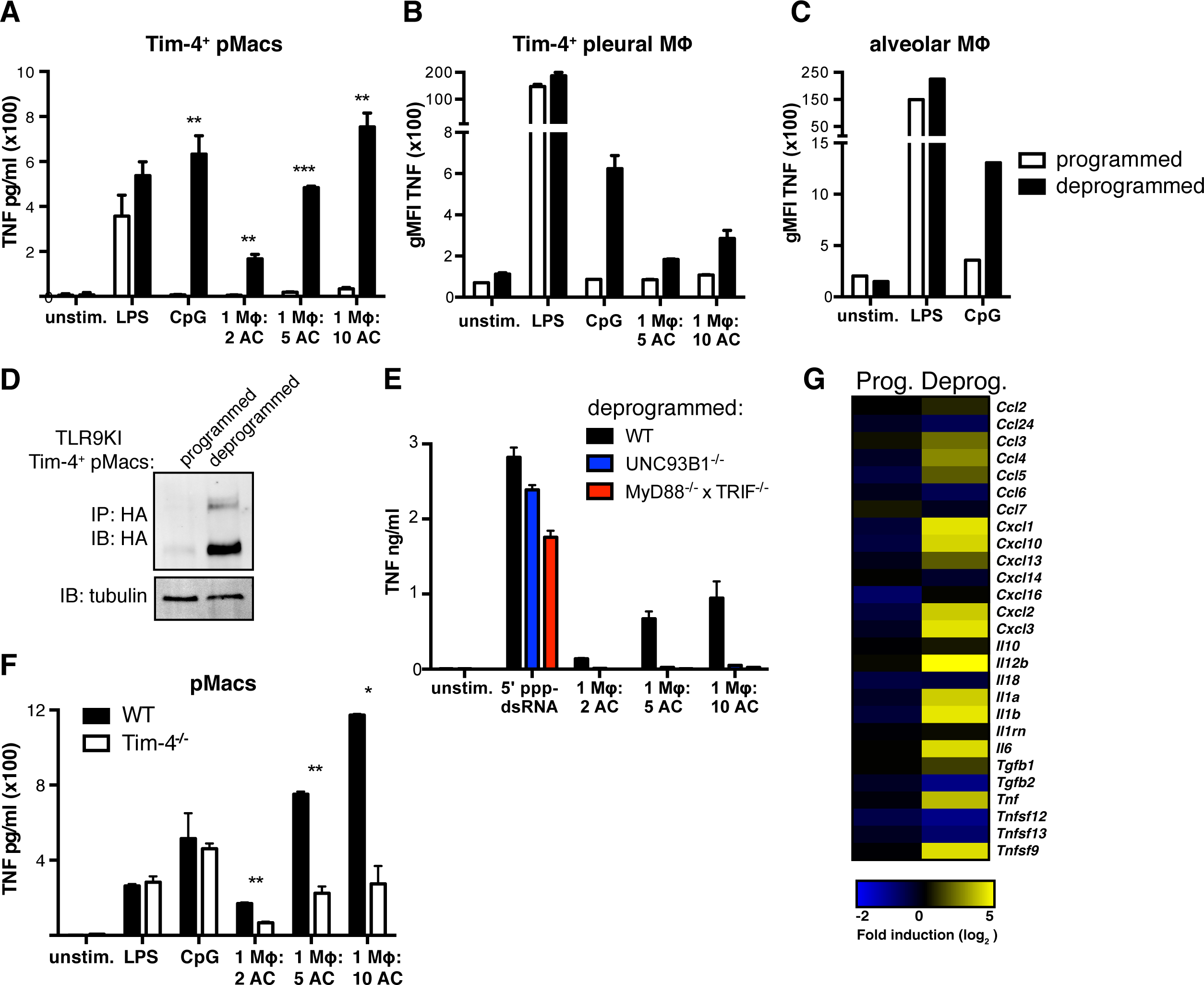
AC-engulfing macrophages removed from their tissue environment generate inflammatory responses to ACs. **(A)** Tim-4^+^ pMacs removed from their tissue environment gain responsiveness to CpG and ACs. Isolated Tim-4^+^ pMacs were cultured overnight (programmed) or for 60 hours (deprogrammed) before stimulation with TLR ligands and ACs. TNF was measured by CBA. Data are representative of at least three independent experiments. **(B)** Tim-4^+^ pleural macrophages removed from their tissue environment gain responsiveness to CpG and ACs. Pleural cells from WT mice were harvested and incubated overnight or for 60 hours before stimulation with TLR ligands and ACs. TNF production was measured by ICS. Data are representative of at least three independent experiments. **(C)** Alveolar macrophages removed from their tissue environment gain responsiveness to CpG. Lung cells from WT mice were harvested and incubated overnight or for 60 hours before stimulation with TLR ligands. TNF production was measured by ICS. Data are representative of at least three independent experiments. **(D)** Tim-4^+^ pMacs removed from their tissue environment gain expression of TLR9. Lysates from programmed and deprogrammed Tim-4^+^ pMacs were immunoprecipitated with anti-HA resin and eluted protein was analyzed by anti-HA western. An anti-tubulin immunoblot was performed on each lysate as a reference. Relative densities normalized to tubulin are 0.036 for programmed and 1 for deprogrammed Tim-4^+^ pMacs. Data are representative of three independent experiments **(E)** Deprogrammed macrophage responses to ACs are dependent on endosomal TLRs. Deprogrammed Tim-4^+^ pMacs from WT, UNC93B1^−/-^, or MyD88^−/-^ TRIF^−/-^ mice were stimulated with TLR ligands and ACs. TNF was measured by CBA. Data are representative of at least three independent experiments. **(F)** Tim-4^−/-^ peritoneal macrophages do not generate robust responses to ACs. Deprogrammed peritoneal macrophages from WT or Tim-4^−/-^ mice were stimulated with TLR ligands and apoptotic cells. Cytokines were measured by CBA. Data are representative of two independent experiments. **(G)** Deprogrammed macrophages upregulate inflammatory cytokine mRNA expression in response to ACs. Programmed and deprogrammed Tim-4^+^ pMacs were stimulated with ACs. RNA was isolated from unstimulated and AC stimulated cells and analyzed by RNA sequencing. Data are presented as fold stimulated over unstimulated and are results from one experiment with technical duplicates averaged so only one value is shown. See also Figure S3 and S4.

To obtain a more complete picture of the responses generated by the *ex vivo* tissue macrophages stimulated with ACs we used RNAseq. Tim-4^+^ pMacs generated a limited transcriptional response to ACs that did not include the production of pro- or anti-inflammatory cytokine mRNA (Figures 3G and S3D). However, *ex vivo* cultured Tim-4^+^ pMacs generated a robust response to ACs, including upregulation of inflammatory cytokines such as *Tnf*, *Il1a*, *Ilb1*, and *Cxcl2* (Figure 3G).

We confirmed that the observed responses were to nucleic acids within ACs rather than free nucleic acids by using Tim-4^−/-^ pMacs. Tim-4 is required for efficient uptake of ACs by pMacs (Miyanishi et al., 2007; Wong et al., 2010) (Figure S3B). Reduced AC uptake by Tim-4^−/-^ pMacs correlated with reduced production of TNF (Figure 3F). Altogether, these results demonstrate that macrophages removed from their tissue of residence can no longer avoid recognition of nucleic acids within ACs. We refer to this change in macrophage responsiveness to ACs as “deprogramming”.

Resident pMacs are influenced by signals from the omentum that induce expression of pMac signature genes (i.e., genes with expression much higher in pMacs relative to their expression in other tissue macrophages). Retinoic acid generated in the omentum induces a number of genes in pMacs, including the transcription factor GATA6, which in turn induces the expression of many other pMac signature genes (Gautier et al., 2014; Okabe and Medzhitov, 2014; Rosas et al., 2014). To test whether these previously described environmental cues were responsible for the AC-clearance program in Tim-4^+^ pMacs, we supplied macrophages undergoing deprogramming with omentum culture supernatant, which has been demonstrated to induce pMac signature genes (Okabe and Medzhitov, 2014). While this treatment improved macrophage survival, it did not prevent the responses to CpG ODN or ACs observed after a three-day culture (Figure S3E, and data not shown). Therefore, for the remainder of these studies we cultured pMacs in omentum culture supernatant during deprogramming to maintain similarity to programmed macrophages as much as possible and to improve their survival *ex vivo*. Next, we directly tested whether GATA6 could induce the AC-clearance program. While deprogrammed macrophages did express reduced levels of GATA6 (Figure S3F), GATA6 overexpression in resident Tim-4^+^ pMacs by *in vivo* lentiviral transduction did not prevent responses to CpG ODN or AC stimulation (Figures S3F and S3G). Therefore, these previously described signals are not sufficient to maintain the AC-clearance program in *ex vivo* tissue macrophages.

To more thoroughly compare programmed and deprogrammed Tim-4^+^ pMacs we analyzed gene expression by RNA sequencing. The expression of many previously identified pMac signature genes decreased during the deprogramming period, further demonstrating that deprogrammed macrophages were losing aspects of their tissue-programmed identity even in the presence of omentum culture supernatant (Figure S4A, (Gautier et al., 2012b)). However, expression of macrophage core genes (i.e., genes expressed by tissue-resident macrophage populations but not dendritic cells) did not show a similar reduction (Figure S4B). In addition, although the expression of some genes previously associated with classical (M1) macrophages increased during the deprogramming period, others decreased. Similarly, expression of some alternatively activated (M2) macrophage associated genes increased while others decreased. Thus, there was no coordinated change in gene expression indicative of a shift from M2 to M1 or vice versa ((Jablonski et al., 2015), Figure S4C). Expression of mRNA for Tim-4 was reduced over ten fold in deprogrammed macrophages; however, protein expression had not decreased enough to affect AC engulfment (Figure S3C and S4D). RNA sequencing results for several of the more relevant genes was confirmed by nCounter analysis of mRNA (Figure S4E).

We also tested directly whether signals from the tissue environment can confer the AC-clearance program. To this end, we examined how gene expression changes when macrophages are transplanted into a tissue. Congenically marked BMMs differentiated with M-CSF for seven days *in vitro* were injected into the peritoneum or administered intranasally. Strikingly, after several weeks in the peritoneum or lungs, a proportion of the transplanted BMMs upregulated Tim-4 or Siglec-F (a marker of alveolar macrophages), respectively (Figures 4A and 4B). Furthermore, when harvested and stimulated with TLR ligands, both populations of transplanted BMMs failed to respond to CpG ODN (Figures 4C and 4D). We excluded the possibility that newly programmed transplanted BMMs arose from a small population of contaminating progenitor cells, since we did not observe significant proliferation of BMMs after IP injection (Figure S4F). Thus, the TLR responses of transferred BMMs were more similar to those of resident macrophages in these tissues than to those of uninjected BMMs.

**Figure 4.**
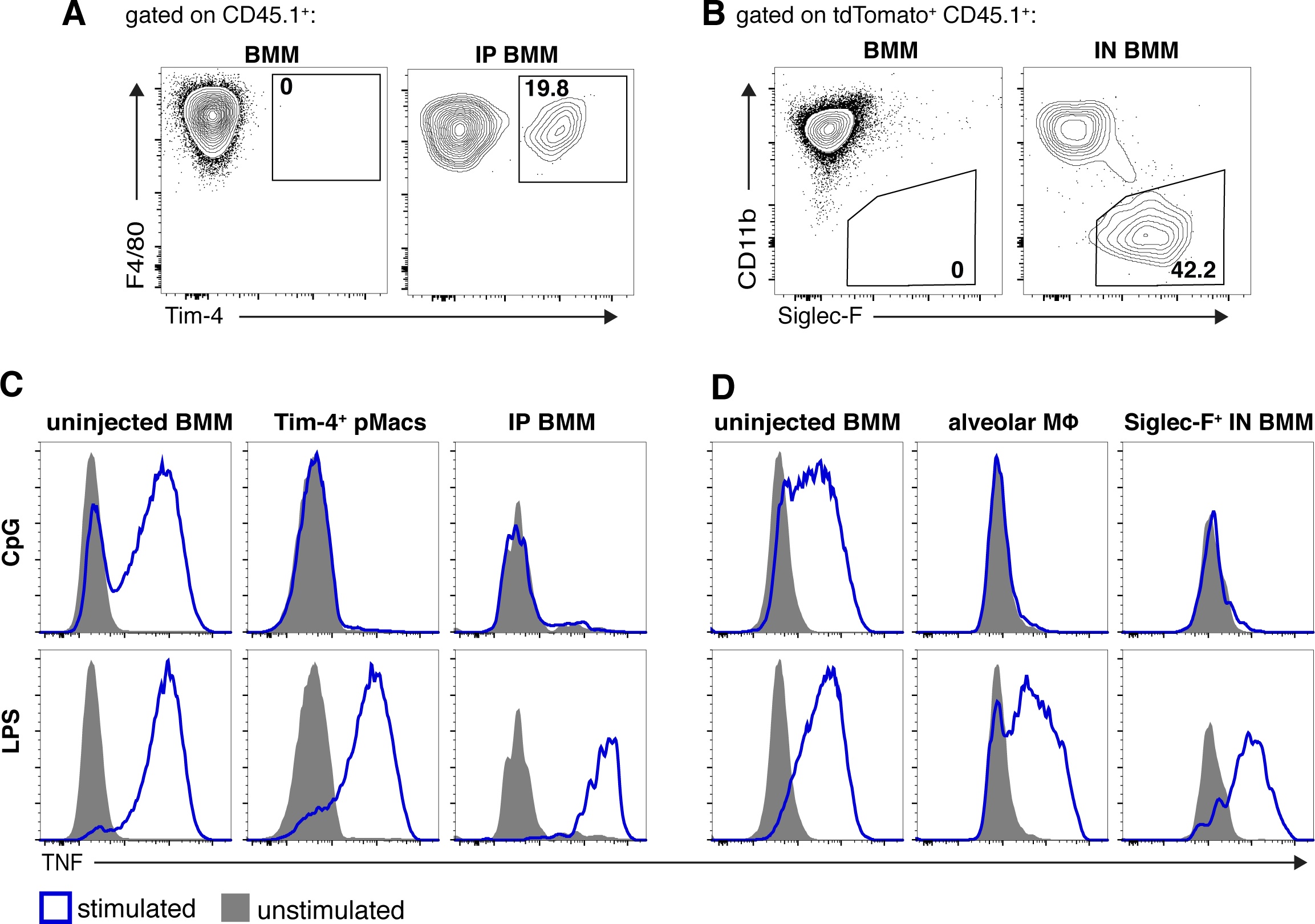
BMMs gain AC-clearance programming in tissue environments. **(A)** In the peritoneal environment BMMs gain Tim-4 expression. After three weeks injected cells were compared to uninjected BMMs. **(B)** In the lung environment BMMs gain Siglec-F expression. After five and a half weeks administered cells were compared to uninjected BMMs. Data are representative of three independent experiments. **(C)** In the peritoneal environment BMMs lose responsiveness to TLR9 ligands. CD45.1^+^WT BMMs were injected into the peritoneum. After three weeks peritoneal cells were harvested and stimulated with TLR ligands. TNF production was measured by ICS. Data are representative of at least three independent experiments with n = 2 or 3 per group per experiment. **(D)** In the lung environment BMMs lose responsiveness to TLR9 ligands. CD45.1 + tdTomato^+^ cells were intranasally (IN) administered. After five and a half weeks lung cells were harvested and stimulated with TLR ligands. Shown are SiglecF^+^ IN administered BMMs. TNF production was measured by ICS. Data are representative of three independent experiments with n = 3-5 per group per experiment See also Figure S4.

These results suggest that environmental cues induce a program in certain populations of tissue macrophages that facilitates silent clearance of ACs. This program induces high expression of AC recognition receptors and low expression of TLR9. Without this programming, the endosomal TLRs of AC-engulfing macrophages robustly recognize and respond to AC-derived nucleic acids.

### Programmed AC-engulfing macrophages have a higher activation threshold for endosomal TLR responses

Next, we more carefully analyzed endosomal TLR signaling in programmed and deprogrammed macrophages. Interestingly, while deprogrammed Tim-4^+^ pMacs macrophages generated a small but consistent phospho-ERK signal in response to ACs, programmed macrophages did not generate this response (Figure 5A). This result indicates that TLR signaling in programmed macrophages is impaired at a step prior to phosphorylation of ERK. We also analyzed gene expression in programmed and deprogrammed Tim-4^+^ pMacs by RNAseq and noted the downregulation of several inhibitors of TLR signaling in deprogrammed macrophages (Figure 5B). Some of these inhibitors, such as SOCS3, DUSP1, and SHP-1, affect TLR signaling prior to ERK phosphorylation (Kondo et al., 2012).

**Figure 5.**
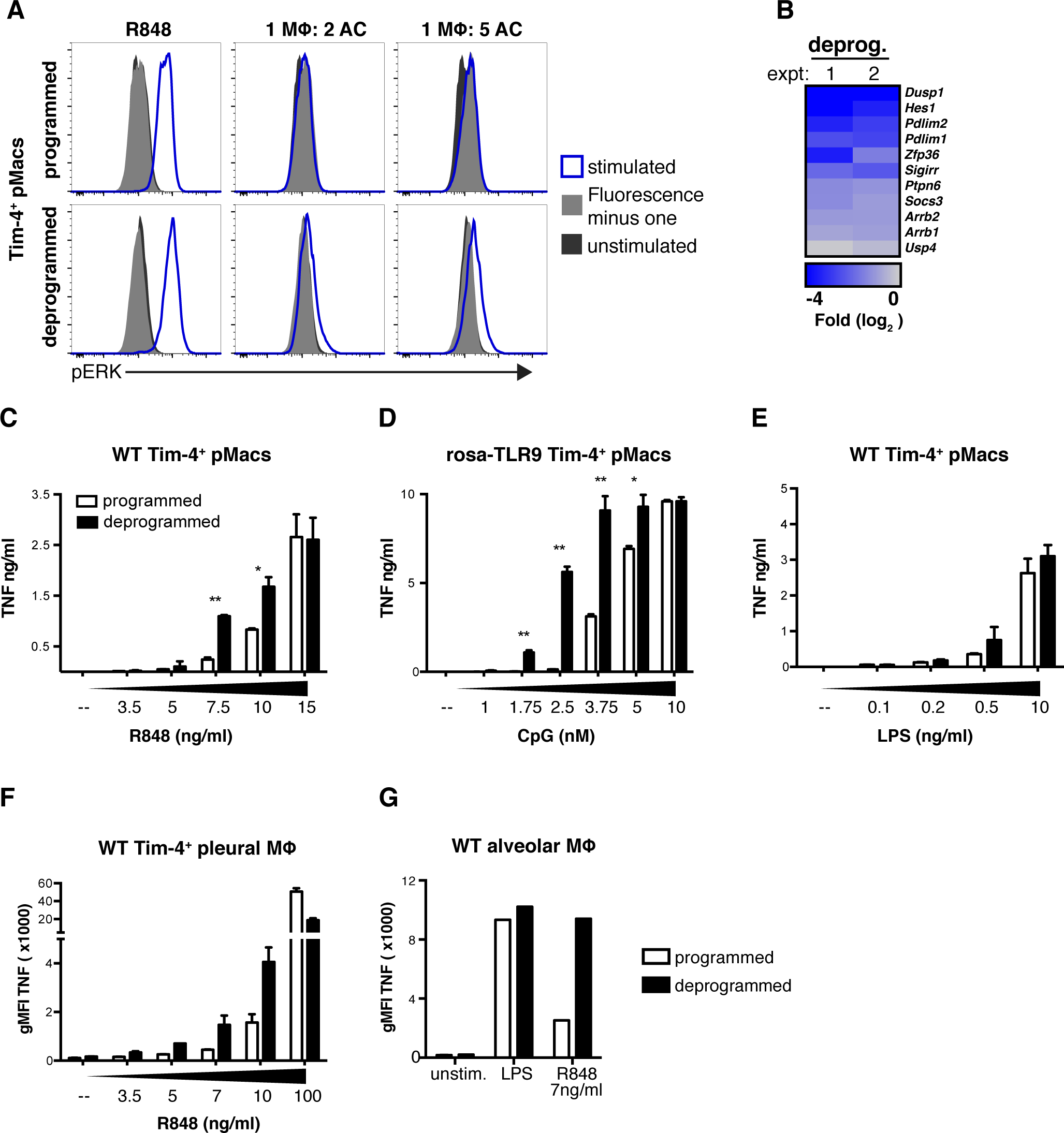
Programmed macrophages have a higher activation threshold for TLR7 and TLR9 responses. **(A)** Deprogrammed macrophages signal in response to ACs while programmed macrophages do not. Programmed and deprogrammed Tim-4^+^ pMacs were stimulated with R848 (1μg/ml) or ACs for 30min, and phospho-ERK1/2 signal was measured by flow cytometry. Data are representative of three independent experiments. **(B)** Deprogrammed Tim-4^+^ pMacs downregulate inhibitors of TLR signaling. RNA was isolated from programmed and deprogrammed WT Tim-4^+^ pMacs and analyzed by RNA sequencing. The heat map indicates genes previously shown to be negative regulators of TLR signaling that are significantly (p < 0.05) downregulated in deprogrammed Tim-4^+^ pMacs relative to programmed Tim-4^+^ pMacs. Results of two independent experiments are shown. Replicates have been averaged so only one value per experiment is shown. **(C)** Deprogrammed Tim-4^+^ pMacs respond to lower doses of TLR7 ligand. Programmed and deprogrammed WT Tim-4^+^ pMacs were stimulated with increasing doses of R848. TNF was measured by CBA. Data are representative of at least three independent experiments. **(D)** Deprogrammed Tim-4^+^ pMacs from rosa-TLR9 mice respond to lower doses of TLR9 ligand. Programmed and deprogrammed Tim-4^+^ pMacs from rosa-TLR9 mice were stimulated with increasing doses of CpG ODN. TNF was measured by CBA. Data are representative of at least three independent experiments. **(E)** Programmed and deprogrammed WT Tim-4^+^ pMacs were stimulated with increasing doses of LPS. TNF was measured by CBA. **(F and G)** Deprogrammed Tim-4^+^ pleural and alveolar macrophages more readily respond to low dose TLR7 ligand. Pleural (F) or lung (G) cells from WT mice were harvested and incubated overnight or for 60 hours before stimulation. TNF production was measured by ICS. Data are representative of two independent experiments. See also Figure S5.

Based on the small signaling response observed in deprogrammed macrophages fed ACs, as well as the reduced expression of several inhibitors of TLR signaling, we hypothesized that the tissue macrophages involved in AC clearance may require a high threshold for initiation of endosomal TLR signaling. This high threshold could prevent responses to weak ligands, such as those derived from ACs. Several aspects of AC-derived nucleic acids make them weak ligands for endosomal TLRs. For instance, mammalian DNA contains very few unmethylated CpG dinucleotides, and upon induction of apoptosis, DNA and RNA are actively degraded into smaller fragments (Coch et al., 2009; Enari et al., 1998; Houge et al., 1993; King et al., 2000; Krieg et al., 1995). To compare the activation threshold for TLR signaling between programmed and deprogrammed macrophages, we measured responses to low doses of TLR ligands. Deprogrammed Tim-4^+^ pMacs, Tim-4^+^ pleural, and alveolar macrophages generated a more robust response to low levels of TLR7 ligands (Figures 5C, 5F, 5G, S5A, and S5B). Importantly, TLR7 expression is not affected by deprogramming (Figure S4D). After deprogramming, Tim-4^+^ pMacs forced to express TLR9 generated significantly enhanced responses to low doses of CpG (Figures 5D and S5C). We observed similar results using CpG ODN with a phosphodiester backbone (PD CpG), which is more labile than the nuclease resistant phosphorothioate CpG ODN (Figure S5D and S5E). Although there were small differences in response to low doses of TLR4 ligand, these were less notable (Figure 5E). Altogether, these data are consistent with the model that macrophages programmed by environmental cues maintain a higher threshold for TLR7 and TLR9 activation and cannot generate responses to the less stimulatory ligands derived from ACs.

### Inflammatory cues induce TLR9 expression without enabling responses to ACs

Although the absence of TLR9 in peritoneal, pleural, and alveolar macrophages may be beneficial when clearing ACs, this lack of expression could be detrimental during infection. We therefore examined how inflammatory signals affect the AC-clearance program in these macrophage populations. As a first step, we incubated purified Tim-4^+^ pMacs overnight with different cytokines followed by stimulation with TLR ligands or ACs. Tim-4^+^ pMacs pretreated with IFNγ or IFNβ upregulated TLR9 expression and gained responsiveness to TLR9 stimulation (Figures 6A, 6B, and S6A). Interestingly, IFNγ or IFNβ treatment did not enable responses to ACs, indicating that the cytokines did not reverse the entire AC-clearance program. This lack of response was not due to an inability to engulf ACs (Figure S6C). IFNγ pretreatment also induced TLR9 responses in Tim-4^+^ pleural and alveolar macrophages (Figure 6C).

**Figure 6.**
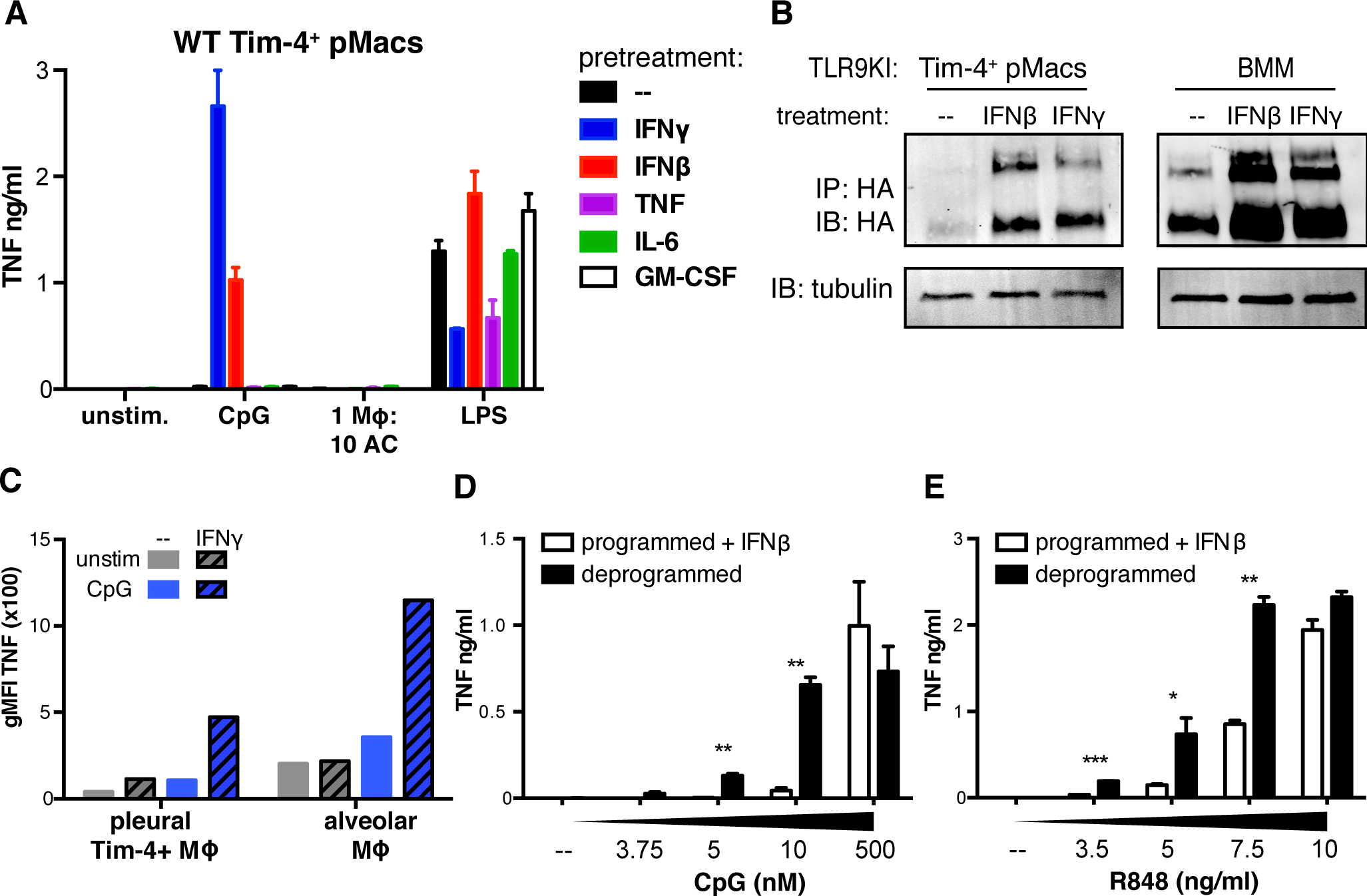
Inflammatory cues induce TLR9 expression in AC-engulfing macrophages. **(A)** IFNγ and IFNβ induce responses to TLR9 ligands in Tim-4^+^ pMacs. Isolated Tim-4^+^ pMacs were treated overnight with the indicated cytokines before stimulation with TLR ligands or ACs. Cytokine responses were measured by CBA. Data are representative of at least three independent experiments. **(B)** IFNγ and IFNβ induce TLR9 expression in Tim-4^+^ pMacs. Tim-4^+^ pMacs and BMMs from TLR9KI mice were cultured overnight ± IFNγ or IFNβ, and TLR9 levels in lysates were measured by anti-HA immunoprecipitation and immunoblot. An anti-tubulin immunoblot was performed on each lysate as a reference. Wells of Tim-4^+^ pMacs correspond to lysate of 1.6e6 cell equivalents, and wells of BMMs correspond to lysate of 0.4e6 cell equivalents. Relative densities normalized to tubulin for Tim-4^+^ pMacs are 1 for untreated, 7.67 for IFNβ treated, and 10.22 for IFNγ treated. Relative densities normalized to tubulin for BMMs are 1 for untreated, 4.50 for IFNβ treated, and 3.37 for IFNγ treated. Data are representative of at least three independent experiments. **(C)** IFNγ induces Tim-4^+^ pleural macrophage and alveolar macrophage responses to TLR9 ligands. Lung and pleural cells from WT mice were harvested and incubated overnight ± 100ng/ml IFNγ before stimulation with CpG ODN. TNF production was measured by ICS. Data are representative of two independent experiments. **(D)** IFNβ treated Tim-4^+^ pMacs maintain a high threshold for TLR9 responses. Isolated programmed IFNβ treated and deprogrammed untreated Tim-4^+^ pMacs were stimulated with increasing doses of CpG. TNF was measured by CBA. Data are representative of two independent experiments. **(E)** IFNβ treated Tim-4^+^ pMacs maintain a high threshold for TLR7 responses. Isolated programmed IFNβ treated and deprogrammed untreated Tim-4^+^ pMacs were stimulated with increasing doses of R848. TNF was measured by CBA. Data are representative of two independent experiments. See also Figure S6.

The lack of response to ACs, despite upregulation of TLR9, in macrophages exposed to inflammatory signals suggested that these cells maintain a high threshold for endosomal TLR signaling. To test this possibility, we compared programmed Tim-4^+^ pMacs that had been pretreated with IFNβ to deprogrammed Tim-4^+^ pMacs. Despite expressing similar amounts of TLR9, IFNβ treated Tim-4^+^ pMacs generated weaker responses to low doses of CpG ODN and R848 when compared to deprogrammed macrophages (Figures 6D, 6E, S6B, S6D, and S6E). These results suggest that IFN treated macrophages maintain a high activation threshold for endosomal TLR signaling, despite upregulation of TLR9. Although prolonged expression of TLR9 in AC-engulfing macrophages may be detrimental, as seen in LysMcre^+/-^Rosa^stop-flox-TLR9^ mice in the pristane-induced lupus model (Figure S2E and S2F), the inability of these macrophages to respond to low-level nucleic acid stimulation may prevent responses to self-derived nucleic acids during acute inflammation.

### KLF2 and KLF4 control an AC-clearance program in macrophages

We reasoned that environmental cues would induce expression of one or more transcription factors that control the AC-clearance program by upregulating TLR inhibitory genes, downregulating TLR9, and upregulating AC recognition receptors. Expression of many transcription factors was lower in deprogrammed Tim-4^+^ pMacs (Figure S7A). These transcription factors included GATA6 and RARβ that have previously been described to influence peritoneal macrophage identity. In addition, the transcription factors Kruppel-like factor 2 (KLF2) and KLF4 were significantly downregulated after deprogramming. These transcription factors contribute to gene regulation in a variety of biological contexts, including vascular biology, cell self-renewal, and, most relevant to our study, macrophage polarization and repression of myeloid cell activation (Chiplunkar et al., 2013; Liao et al., 2011; Mahabeleshwar et al., 2011; Soucie et al., 2016).

To examine whether these transcription factors contribute to the AC-clearance program we first assessed their importance for inhibition of TLR9 expression in the peritoneal environment. To this end, we used Cas9 genome editing to generate BMMs lacking each transcription factor (Figure S7B). These BMMs were injected into the peritoneums of congenically marked mice. Despite low recovery of injected BMMs, we could demonstrate that after two weeks *in vivo* the BMMs transduced with empty vector or with guides for GATA6 or RARβ lost the ability to respond to TLR9 stimulation, similar to what we observed previously for transferred untransduced BMMs (Figures 7A and 7B). In contrast, BMMs expressing guides specific for KLF2 continued to respond to TLR9 stimulation, and KLF4 targeted BMMs had an intermediate response (Figures 7A and 7B). These data suggest that KLF2, potentially with contributions from KLF4, inhibits TLR9 responses in resident pMacs.

**Figure 7.**
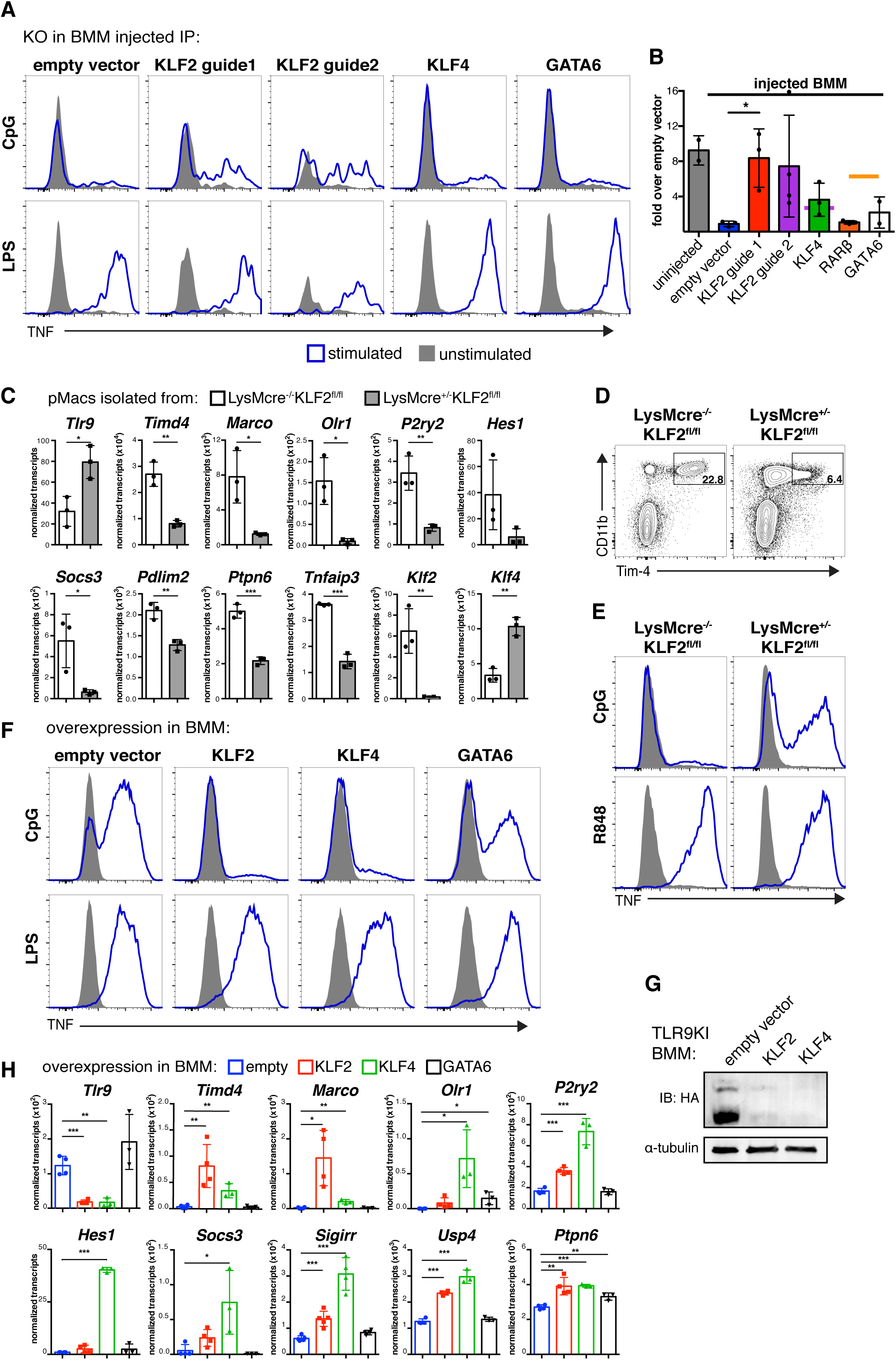
KLF2 and KLF4 imprint an AC-clearance program on macrophages. **(A and B)** In the peritoneal environment, BMMs lacking KLF2 maintain responses to TLR9 stimulation. BMMs from cas9 overexpressing mice were transduced with vectors encoding guide RNAs targeting the indicated transcription factors or with empty vector. After differentiation *in vitro*, BMMs were injected intraperitoneally into congenically marked mice. 14 days after injection peritoneal cells were harvested and stimulated with TLR ligands. Uninjected BMMs transduced with empty vector were stimulated as a control. **(B)** TNF production was measured by ICS. Data are representative of at least two independent experiments. (C) Quantification of IP injected BMMs responses to CpG displayed relative to injected empty vector response. Data are the combined results from two independent experiments with total n = 2-4 per group. **(C)** Expression of AC recognition receptors and negative regulators of TLR signaling is reduced in pMacs lacking KLF2. F4/80^+^ pMacs were isolated from LysMcre^+/-^KLF2^fl/fl^ mice and littermate control LysMcre^−/-^KLF2^fl/fl^ mice. The levels of mRNA for the indicated genes were quantified using a Nanostring nCounter. n = 3 per group from independent experiments. **(D and E)** pMacs lacking KLF2 demonstrate reduced expression of Tim-4 and are responsive to TLR9 ligands. Peritoneal cells from LysMcre^+/-^KLF2^fl/fl^ mice and littermate control LysMcre^−/-^KLF2^fl/fl^ mice were harvested and stimulated with TLR ligands. (D) Unstimulated cells were analyzed for expression of Tim-4. Data are representative of three independent experiments. **(F)** Overexpression of KLF2 or KLF4 in BMMs dampens responses to CpG. WT BMMs were transduced with retroviral vectors encoding KLF2, KLF4, GATA6 or with empty vector. Cells were stimulated with TLR ligands and TNF production was measured by ICS. Data are representative of at least two independent experiments. **(G)** Overexpression of KLF2 and KLF4 in BMMs prevents expression of TLR9. TLR9KI BMMs were transduced with retroviral vectors encoding KLF2, KLF4, or empty vector. Lysates were subjected to immunoprecipitation with anti-HA resin and eluted TLR9 protein was visualized by anti-HA immunoblot. An anti-tubulin immunoblot on the same lysates was performed as a reference. Relative densities normalized to tubulin are 1 for empty vector, 0.06 for KLF2, and 0.02 for KFL4. Data are representative of at least two independent experiments. **(H)** Overexpression of KLF2 and KLF4 in BMMs induces expression of AC recognition receptors and negative regulators of TLR signaling. BMMs were transduced with retroviral vectors encoding KLF2, KLF4, GATA6 or with empty vector. The levels of mRNA for the indicated genes were quantified using a Nanostring nCounter. n = 3-4 per group from independent experiments. See also Figure S7.

To determine whether KLF2 controls expression of other aspects of the AC-clearance program, we analyzed gene expression in pMacs isolated from LysMcre^+/-^ KLF2^fl/fl^ mice and LysMcre^−/-^KLF2^fl/fl^ littermate control mice. Expression of Tim-4 was reduced in KLF2-deficient cells, as was expression of the AC recognition receptors Marco and Olr1 and *P2ry2*, an ATP receptor demonstrated to aid in recruitment of phagocytes to ACs (Figure 7C) (Elliott et al., 2009; Oka et al., 1998; Wermeling et al., 2007). We confirmed reduced expression of Tim on the surface of LysMcre^+/-^KLF2 ^fl/fl^ pMacs by flow cytometry (Figure 7D). Remarkably, high expression of negative regulators of TLR signaling also required KLF2; mRNA levels of Hes1, Socs3, Pdlim2, Ptpn6, and Tnfaip3 KLF2-deficient pMacs. LysMcre^+/-^KLF2 ^l/fl^ pMacs also expressed higher levels of TLR9 than littermate controls and responded to CpG ODN (Figure 7E). These results indicate that KLF2 is required for expression of all aspects of the AC-clearance program.

We also tested whether KLF2 and KLF4 are sufficient to drive expression of the AC-clearance program in BMMs. KLF2- and KLF4-overexpressing BMMs showed dramatically reduced TLR9 responses but not TLR4 responses, while GATA6 overexpression had little effect (Figures 7F, S7C, and S7D). KLF2 and KLF4 inhibited TLR9 expression and drove expression of many of the AC-clearance program genes shown to be KLF2-dependent from our analysis of LysMcre^+/-^KLF2^fl/fl^ pMacs (Figures 7G and 7H).

These data demonstrate that the transcription factors KLF2 and KLF4 control a program in macrophages that facilitates silent clearance of ACs. These transcription factors induce a series of coordinated changes, including the downregulation of TLR9 and the upregulation of inhibitors of TLR signaling and receptors that recognize ACs. Notably, while alveolar macrophages express low levels of KLF2, their expression of KLF4 is quite high (data from the Immunological Genome Project), suggesting that KLF4 may play a more prominent role in programming alveolar macrophages while KLF2 plays a similar role in pMacs.

## DISCUSSION

The characteristics that enable macrophages to clear ACs in an immunologically silent manner remain unclear. Recent studies demonstrating the heterogeneity of tissue-resident macrophages highlight the functional differences between populations of macrophages. Here we have characterized three macrophage populations adept at AC clearance in the peritoneum, pleural cavity, and lungs. We demonstrate that these AC-engulfing macrophages are programmed by their tissue environments to clear ACs in an immunologically silent manner. This programming includes inhibition of TLR9 expression as well as upregulation of AC recognition receptors and negative regulators of TLR signaling. We hypothesize that the resulting increased threshold of activation in these cells enables immunologically silent clearance of large numbers of ACs.

Several mechanisms that limit inflammatory signaling in cells that eat ACs have been previously described, many involving induction of inhibitors upon AC recognition, but our work identifies distinct features intrinsic to certain tissue macrophages that enable silent AC clearance. A recent study examining phagocyte populations that engulf apoptotic epithelial cells in the intestine identified two macrophage populations and a DC population (Cummings et al., 2016). After AC engulfment these populations demonstrated changes in gene expression profiles; all three populations downregulated genes involved in inflammatory responses and upregulated negative regulators of the immune response. In contrast, the macrophages we have characterized appear to be preprogrammed for silent AC clearance and do not demonstrate large changes in gene expression after engulfment of ACs (Figure S3D). We also did not observe upregulation of anti-inflammatory cytokines in pMacs in response to ACs (Figure 3G). The Tyro3, Axl, and Mer (TAM) receptors recognize ACs and have been demonstrated to inhibit TLR signaling by upregulating expression of SOCS genes (Rothlin et al., 2007). However, expression of TAM receptors did not decrease in deprogrammed macrophages, and SOCS genes were not upregulated in programmed macrophages in response to ACs (data not shown). Therefore, although we cannot rule out a role for TAM receptor signaling *in vivo*, we do not believe that the AC-clearance program present in peritoneal, pleural, and alveolar macrophages involves this receptor family.

Interestingly, the three AC-engulfing macrophage populations we have identified reside in cavities and are motile. Many other tissue-resident macrophage populations are immobilized (Stamatiades et al., 2016). A recent report demonstrated that peritoneal macrophages can be recruited directly from the peritoneal cavity into the liver to repair damage after liver injury (Wang and Kubes, 2016). It has also been shown that ACs release find-me signals that recruit phagocytes to dying cells (Elliott et al., 2009; Truman et al., 2008). Perhaps these macrophages are particularly adapted to engulf ACs since they can be recruited to dying cells in their cavities as well as neighboring organs. As cavity resident cells these macrophage populations may also have limited exposure to circulating proteins, such as DNASE1L3, which digest DNA in microparticles released from ACs (Sisirak et al., 2016), One interesting possibility is that cavity resident macrophages utilize the additional mechanisms we describe to limit responses to ACs because of reduced DNase activity in these compartments.

We identify KLF2 and KLF4 as transcription factors critical for induction of an AC-clearance program in macrophages (Figure 7). We demonstrate that KLF2 and KLF4 induce expression of AC recognition receptors as well as negative regulators of TLR signaling. In this way these transcription factors establish an AC-clearance program that links enhanced AC-engulfment ability with reduced TLR signaling. By linking these two features, KLF2 and KLF4 may ensure that macrophages with enhanced abilities to engulf ACs do not generate inappropriate responses to self nucleic acids. A previous study demonstrated that mice lacking KLF2 in LysM^+^ cells display increased levels of proinflammatory cytokines at steady state in their sera and show enhanced inflammation during infection (Mahabeleshwar et al., 2011). Perhaps dysregulation of macrophage AC engulfment ability and endosomal TLR responses contributes to this inflammatory phenotype.

KLF2 and KLF4 expression is not restricted to AC-engulfing macrophages (data from the Immunological Genome Project). Therefore, it is likely that other tissue-induced transcription factors and differences in chromatin structure are involved in induction of the AC-clearance program. Although some transcription factors involved in environmental programming are restricted to a single macrophage population (e.g., GATA6 in peritoneal macrophages) others have more complex expression profiles. For instance the transcription factor PPARγ is required for the development and identity of alveolar macrophages as well as splenic red pulp macrophages (Gautier et al., 2012a; Schneider et al., 2014).

It remains to be determined how tissue environments impart an AC-clearance program. A recent study demonstrating that KLF4 expression is upregulated after macrophage precursors colonize the lung and differentiate into alveolar macrophages, suggests that KLF4 expression is induced by the lung microenvironment (Mass et al., 2016). We were unable to demonstrate a role for the recognition of ACs as the initiating signal (data not shown). However, it may be difficult to recapitulate *in vitro* the continual engulfment of ACs that occurs *in vivo*. It remains possible that continual recognition of ACs could program AC-engulfing macrophages. It is interesting to note that KLF2 and KLF4 expression is induced in endothelial cells following exposure to extracellular ATP released in response to shear stress (Sathanoori et al., 2015). ATP is also a known find-me signal released by ACs that induces phagocyte recruitment (Elliott et al., 2009). Continual exposure of AC-engulfing macrophages to ATP may induce expression of KLF2 and KLF4. We demonstrated that KLF2 and KLF4 expression upregulates P2ry2, the receptor that recognizes this find-me signal, suggesting there could be feed forward regulation of ATP detection and expression of KLF2 and KLF4 (Figure 7F).

Programmed AC-engulfing macrophages have a higher activation threshold for endosomal TLR signaling. This feature likely prevents responses to less stimulatory self ligands, while allowing responses to microbial nucleic acids. Endosomally localized negative regulators of TLR signaling may regulate this threshold. It is also possible that trafficking of endosomal cargo is different in programmed versus deprogrammed macrophages. LC3-associated phagocytosis (LAP) has previously been demonstrated to facilitate phagosome maturation and degradation of internalized pathogens and ACs (Martinez et al., 2011; Sanjuan et al., 2007). Mice with deficiencies in LAP components demonstrate defective clearance of ACs and develop an autoinflammatory disorder (Martinez et al., 2016). Differences in the ability of deprogrammed macrophages to induce LAP may enable responses to AC-derived cargo.

Expression of certain endosomal TLRs in human myeloid cells is thought to be more limited than their expression in mice. In human peripheral blood mononuclear cells, TLR7 and TLR9 are exclusively expressed in pDCs, while TLR8 is more broadly expressed in macrophages and conventional DCs (Kadowaki et al., 2001; Krug et al., 2001). These data have often been cited to support the conclusion that no human macrophages express these endosomal TLRs. However, given our growing appreciation of tissue macrophage heterogeneity, this concept is worth revisiting. Assuming a similar degree of heterogeneity is present in humans as is seen in mice, it may be interesting to examine TLR expression in different human macrophage populations. For instance, human decidual macrophages have been shown to express TLR3, TLR7, TLR8 and TLR9 in addition to TLR2 and TLR4 (Duriez et al., 2014). It will be important to expand our understanding of nucleic acid sensing TLRs in human tissue-resident macrophages with different functions, especially those that adeptly engulf ACs.

Finally, our work suggests that understanding how the expression of TLRs and regulators of TLR signaling is controlled in specialized cell types may have therapeutic value. We have demonstrated that expression of TLR9 in AC-engulfing macrophages can be modulated by inflammatory signals such as IFN**β** and IFN**γ** (Figure 6). While this induction may enable pathogen detection during infections, we also observe that forced upregulation of TLR9 subtly exacerbates inflammation during a chronic inflammatory disease (Figure S2). An IFN-signature has been described in subsets of patients suffering from autoimmune disorders, most notably lupus (Bennett et al., 2003). The link between type I IFN and disease in these patients is clearly complex, but one mechanism may be upregulation of TLR expression in certain cell types. For example, IFN exposure may upregulate expression of nucleic acid sensing TLRs in macrophages that clear ACs, and this continual expression may promote and prolong inflammation. If human AC-engulfing macrophages also utilize KLF2 and KLF4 to limit expression of endosomal TLRs and promote expression of negative regulators of TLR signaling, then increasing KLF2 or KLF4 expression may reduce inflammatory signaling. In this regard, it is notable that statin treatment can upregulate expression of KLF2 in human peripheral blood monocyte-derived macrophages (Tuomisto et al., 2008). Defining the gene expression programs and unique functional characteristics of distinct tissue macrophage populations will likely reveal additional therapeutic targets and open new avenues for treatment of immune disorders.

## Author Contributions

A.W.R. and G.M.B designed experiments. A.W.R. performed experiments. B.L.L. generated the TLR9 reporter mouse. J.D. analyzed RNA sequencing data. S.J. and M.J.S. generated the Rosa^stop-flox-TLR9^ mouse. A.W.R. and G.M.B. wrote the manuscript.

## Acknowledgements

We thank Dr. Jerry Lingrel for providing LysMcre^+/-^KLF2^fl/fl^ mice. We thank K. Ching for assistance with cloning and R. Vance and members of the Barton and Vance labs for constructive discussions and critical reading of the manuscript. This work was supported by the NIH (AI072429, AI105184 and AI063302 to G.M.B. and AI118841 and AR050256 to M.J.S) and by the Lupus Research Institute (Distinguished Innovator Award to G.M.B.). J.D. was supported by a Long-Term Human Frontier Science Program Fellowship (LT-000081-2013L).

## Supplemental Figure Legends

**Figure S1.**
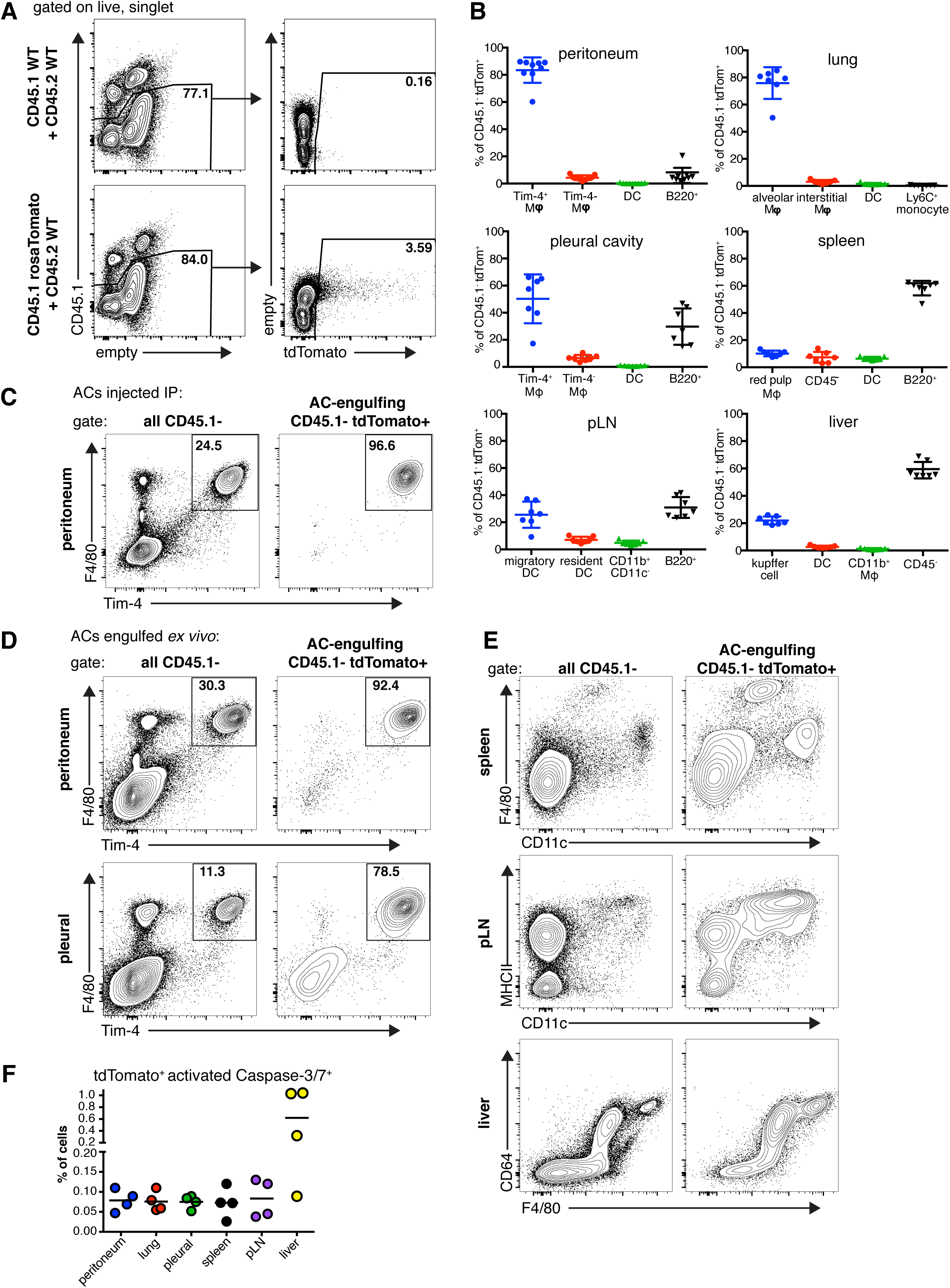
Identification of tissue resident macrophage populations that clear apoptotic cells *in vivo*, related to Figure 1. **(A)** Example gating strategy to identify AC-engulfing cells. Control bone marrow chimeras generated with WT CD45.1^+^ bone marrow were compared to chimeras generated with tdTomato^+^CD45.1^+^ bone marrow. **(B)** Quantification of AC-engulfing (tdTomato+CD45.1 -) cells in different tissues. Data from Figure 1D is represented relative to other cells in the same tissue. Data are the combined results from three independent experiments with total n = 7-9 for all groups. **(C)** Tim-4^+^ macrophages engulf injected ACs. One hour after CD45.1^−^tdTomato^+^ACs were injected intraperitoneally peritoneal cells were harvested and analyzed by flow cytometry. **(D)** Tim-4^+^ peritoneal and pleural macrophages engulf ACs *ex vivo*. All cells from peritoneal or pleural lavage were plated *ex vivo* and incubated with CD45.1^−^ tdTomato^+^ACs before analysis by flow cytometry. **(E)** Representative flow cytometric analysis of tissues from bone marrow chimeras. Cells were gated on CD45.1^−^ cells or AC-engulfing cells (CD45.1^−^tdTomato^+^), as indicated. Data are representative of three independent experiments. **(F)** Quantification of tdTomato^+^ ACs in different tissues. Cells from bone marrow chimeras were assayed for Caspase-3/7 activation. Shown is the percentage of total cells that were tdTomato^+^ and positive for activated Caspase -3/7. Data are the combined results of two independent experiments.

**Figure S2.**
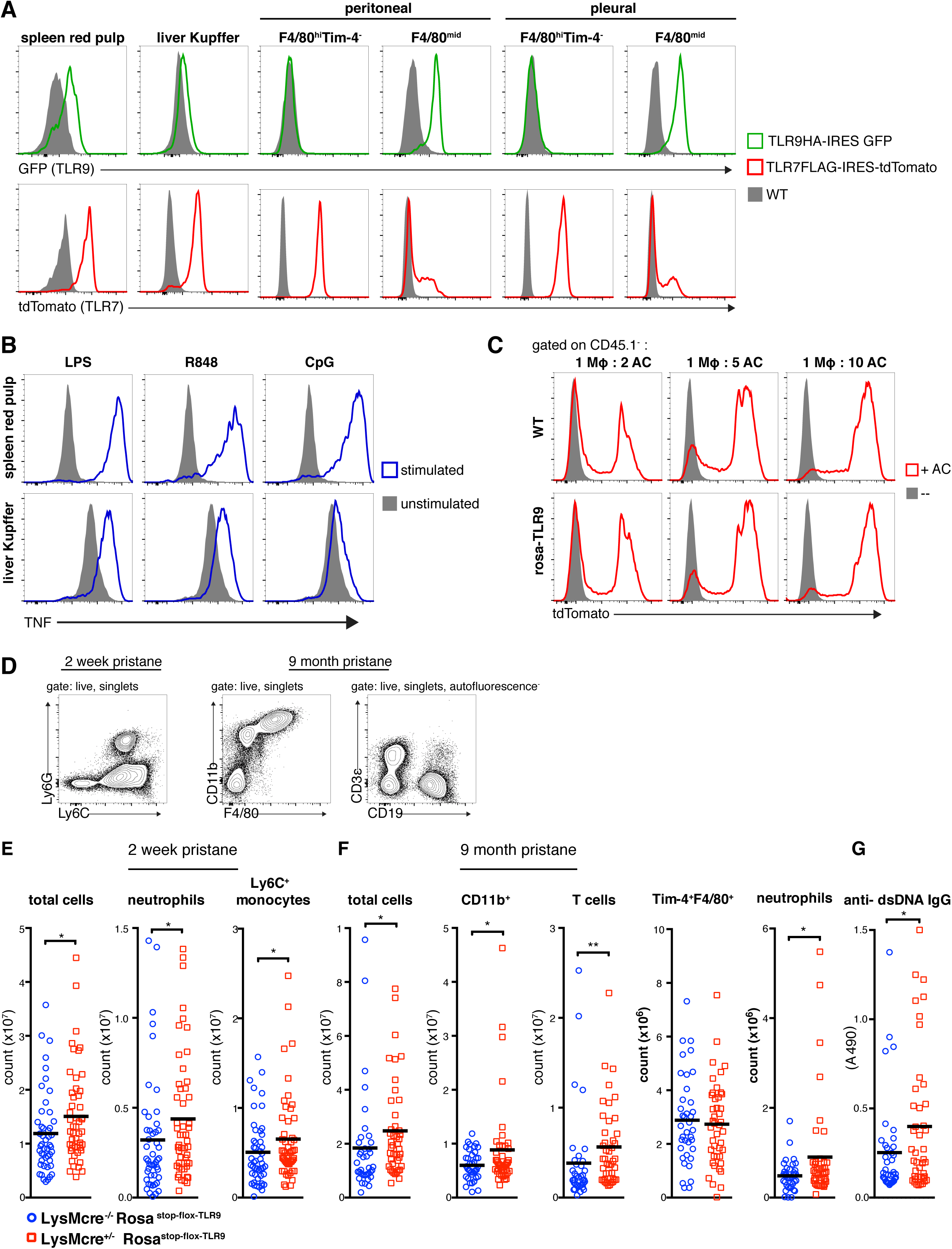
Red pulp macrophages, Kupffer cells, and F/480^mid^ peritoneal and pleural macrophages express TLR9, related to Figure 2. **(A)** Splenic red pulp macrophages and liver Kupffer cells express TLR9 and TLR7. Representative histograms of TLR expression by gated macrophages harvested from TLR9KI, TLR7KI, or WT mice. Red pulp macrophages are gated as live F4/80hiCD11 b^−^ cells. Kupffer cells are gated as live CD45^+^CD64^+^MHCII^+^ cells. F4/80^mid^ macrophages are gated as live CD11b+F4/80 ^mid^ MHCII^+^Ly6C^−^CD11 c^−^. Data are representative of at least three independent experiments with n = 2 per experiment. **(B)** Splenic red pulp macrophages and liver Kupffer cells respond to TL9 ligands. Macrophages from WT mice were stimulated *ex vivo* with TLR ligands. TNF production was measured by ICS. Data are representative of at least three independent experiments with n = 2 per experiment. **(C)** Rosa-TLR9 macrophages are adept at AC engulfment. Tim-4^+^ pMacs were isolated from WT and rosa-TLR9 mice. Macrophages were cultured overnight then incubated with CD45.1^+^tdTomato^+^ ACs at indicated ratios and analyzed by flow cytometry. **(D)** Representative flow cytometric analysis of peritoneal cells at different time points after pristane injection. **(E)** Mice overexpressing TLR9 in LysM^+^cells have slightly enhanced cell recruitment 14 days after pristane injection. LysMcre^+/-^Rosa^stop-flox-TLR9^ or littermate control LysMcre^−/-^Rosa^stop-flox-TLR9^ mice were injected intraperitoneally with pristane. 14 days after injection peritoneal cells were harvested and analyzed by flow cytometry. The number of total peritoneal exudate cells,CD11b^+^ cells, T cells (CD3_ε_^+^CD11b-CD19-), Tim-4+ macrophages neutrophils (CD11b^+^Ly6G^+^Ly6C^mid^), and Ly6C^+^ monocytes (CD11b^+^Ly6C^+^Ly6G^−^) are shown. Data are the combined results of five independent experiments with total n = 48 for LysMcre^−/-^ and n = 51 for LysMcre^+/-^; p-value determined by t-test performed on log-transformed data to account for the non-normal distribution of the data. **(F and G)** Mice overexpressing TLR9 in LysM^+^ cells have slightly enhanced cell counts and anti-dsDNA IgG 9 months after pristane injection. LysMcre^+/-^Rosa^stop-flox-TLR9^ or littermate control LysMcre^−/-^Rosa^stop-flox-TLR9^ mice were injected intraperitoneally with pristane and analyzed 9 months after injection. (D) Peritoneal cells were harvested and analyzed by flow cytometry. The number of total peritoneal exudate cells, CD11b^+^ cells, T cells (CD3ε^+^CD11b^−^CD19^−^), Tim-4^+^ macrophages, and neutrophils (CD11b^+^Ly6G^+^Ly6C^mid^) are shown. (E) Serum samples were collected and tested by ELISA for anti-dsDNA IgG. Data are the combined results of four independent experiments with total n = 38 for LysMcre^−/-^ and n = 43 for LysMcre^+/-^; p-value determined by t-test performed on log-transformed data to account for the non-normal distribution of the data.

**Figure S3.**
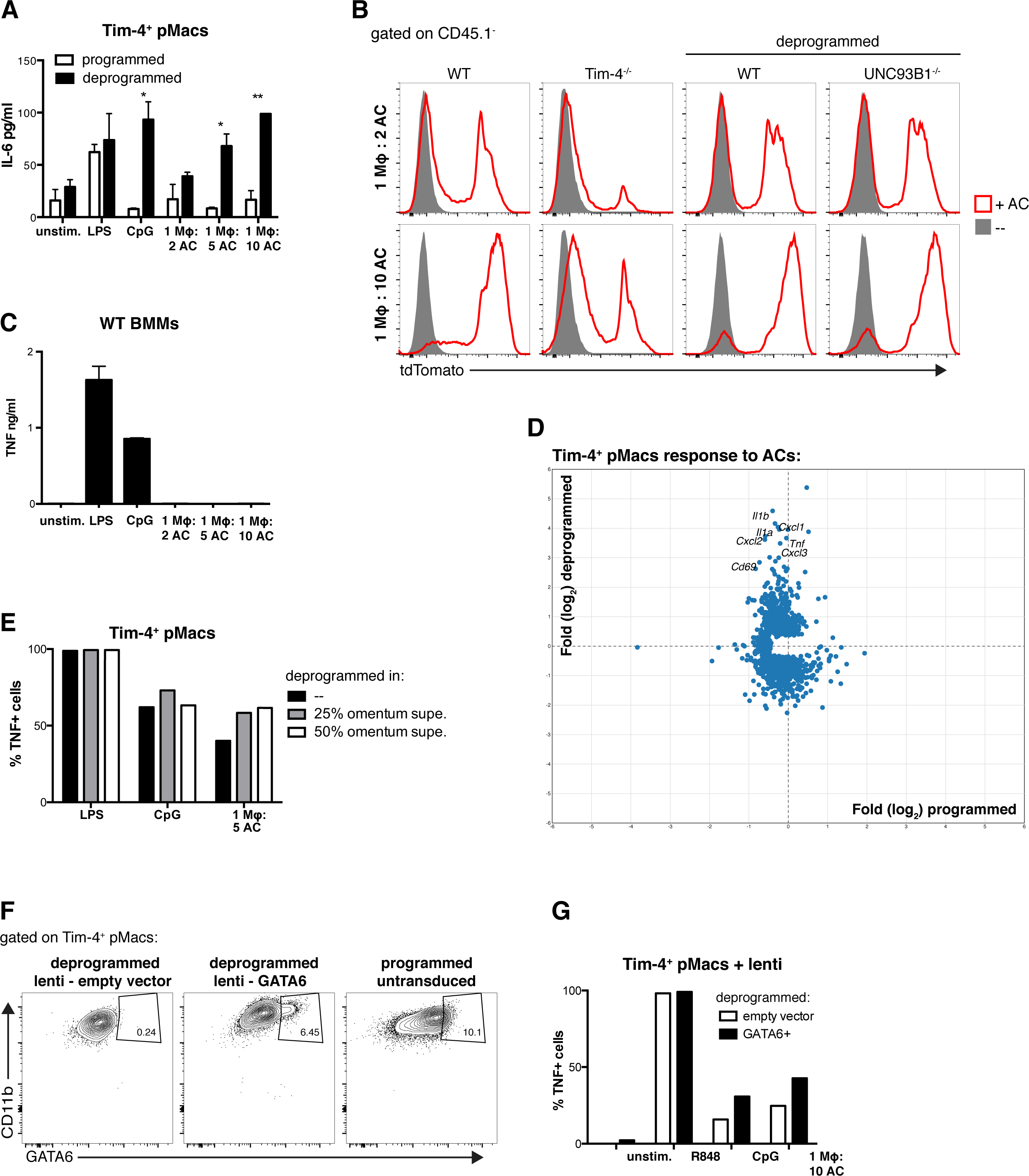
AC-engulfing macrophages removed from their tissue environment generate inflammatory responses to ACs, related to Figure 3. **(A and B)** Tim-4^+^ pMacs removed from their tissue environment gain responsiveness to CpG and ACs. Isolated Tim-4^+^ pMacs were cultured overnight (programmed) or for 60 hours (deprogrammed) before stimulation with TLR ligands and ACs. IL-6 and TNF were measured by CBA. Data are representative of at least three independent experiments. PD CpG - phosphodiester CpG ODN. **(C)** Deprogrammed macrophages are adept at AC engulfment. Isolated Tim-4^+^ pMacs were cultured overnight or for 60 hours then incubated with CD45.1^+^tdTomato^+^ ACs at indicated ratios and analyzed by flow cytometry. **(D)** Deprogrammed Tim-4^+^ pMacs upregulate inflammatory cytokine mRNA in response to ACs while programmed Tim-4^+^ pMacs generate a limited transcriptional response. RNA was isolated from unstimulated and AC stimulated macrophages and analyzed by RNA sequencing. Data are presented as fold stimulated over unstimulated of all significantly changed genes plotted as log2 fold changes. To minimize contamination from AC mRNA, genes whose expression was greater than two fold higher in RNA isolated from ACs alone relative to macrophage RNA were considered AC-derived and excluded. **(E)** BMMs do not generate responses tp ACs. WT BMMs were stimulated with TLR ligands and ACs. TNF were measured by CBA. Data are representative of at least three independent experiments. **(F)** Omentum-derived factors do not prevent deprogramming of pMacs. Peritoneal cells were cultured in complete media alone or complete media plus 25% or 50% omentum supernatant for 60 hours before stimulation with TLR ligands or ACs. TNF production was measured by ICS. Data are representative of at least three independent experiments. **(G and H)** GATA6 overexpression does not prevent deprogramming of pMacs. Mice were injected intraperitoneally with lentivirus expressing GATA6 or empty vector. One week after injection peritoneal cells were harvested and cultured for 60hrs before stimulation with TLR ligands or ACs. (G) GATA6 expression was compared to directly *ex vivo* pMacs. (H) TNF production was measured by ICS. Data are representative of two independent experiments

**Figure S4.**
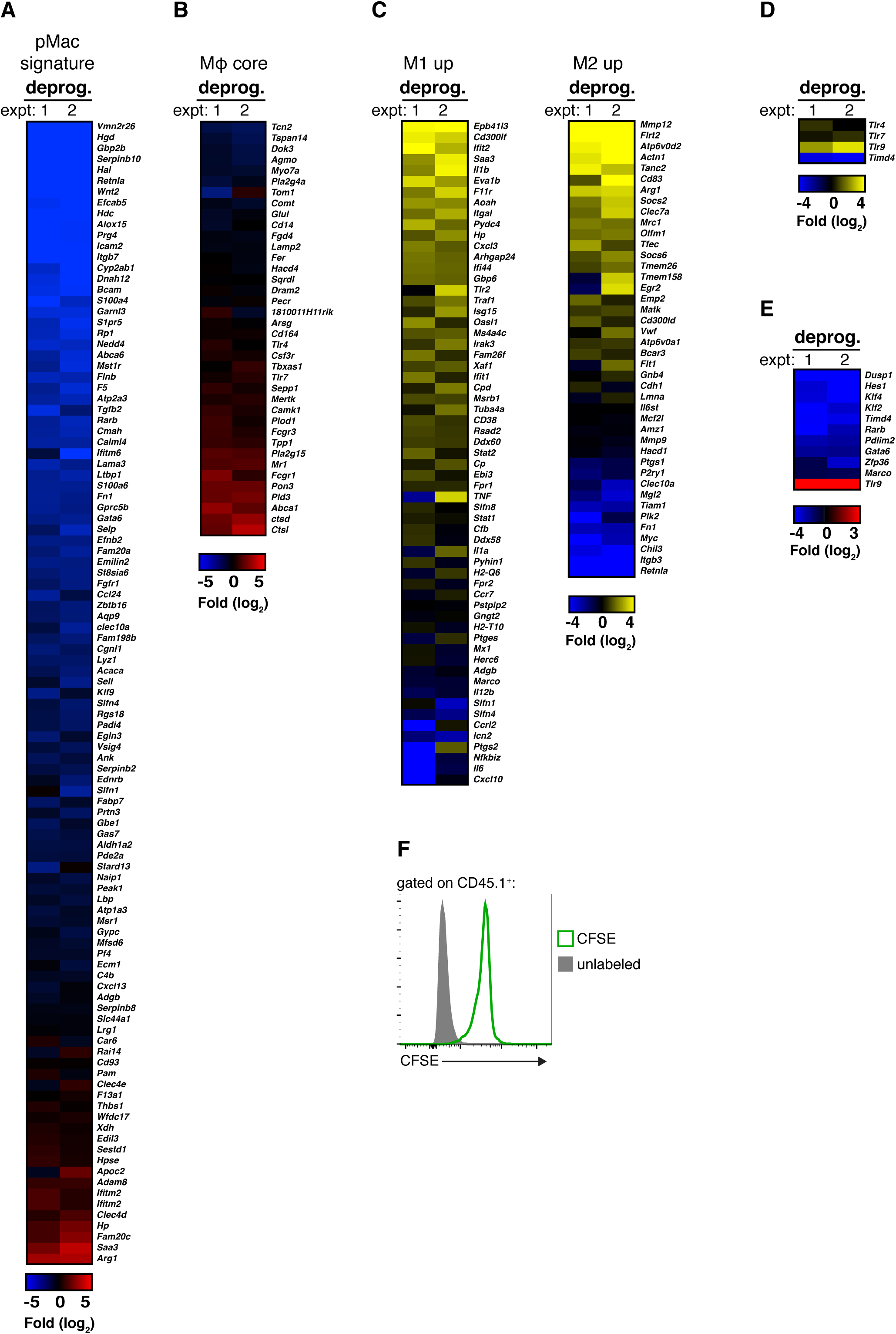
BMMs gain AC-clearance programming in tissue environments related to Figures 3 and 4. **(A - D**) RNA sequencing results from two independent experiments are represented as heat maps of expression of the indicated genes in deprogrammed relative to programmed Tim-4^+^ pMacs. Results from replicates are averaged so only one value per experiment is shown **(A)** Deprogrammed Tim-4^+^ pMacs downregulate many peritoneal macrophage signature genes **(B)** Deprogrammed Tim-4^+^ pMacs do not downregulate macrophage core genes. **(C)** Deprogrammed Tim-4^+^ pMacs expression of classical (M1) and alternative (M2) macrophage associated genes. **(D)** Deprogrammed Tim-4^+^ pMacs upregulate *Tlr9* and downregulate *Timd4*. **(E)** RNA sequencing results for listed genes were confirmed using a Nanostring nCounter. Results from two independent experiments are represented as heat maps of expression of the indicated genes in deprogrammed relative to programmed Tim-4^+^ pMacs. **(F)** IP injected BMMs do not undergo notable proliferation. CD45.1 + BMMs were labeled ± CFSE and injected IP. After three weeks peritoneal cells were harvested and analyzed for CFSE levels.

**Figure S5.**
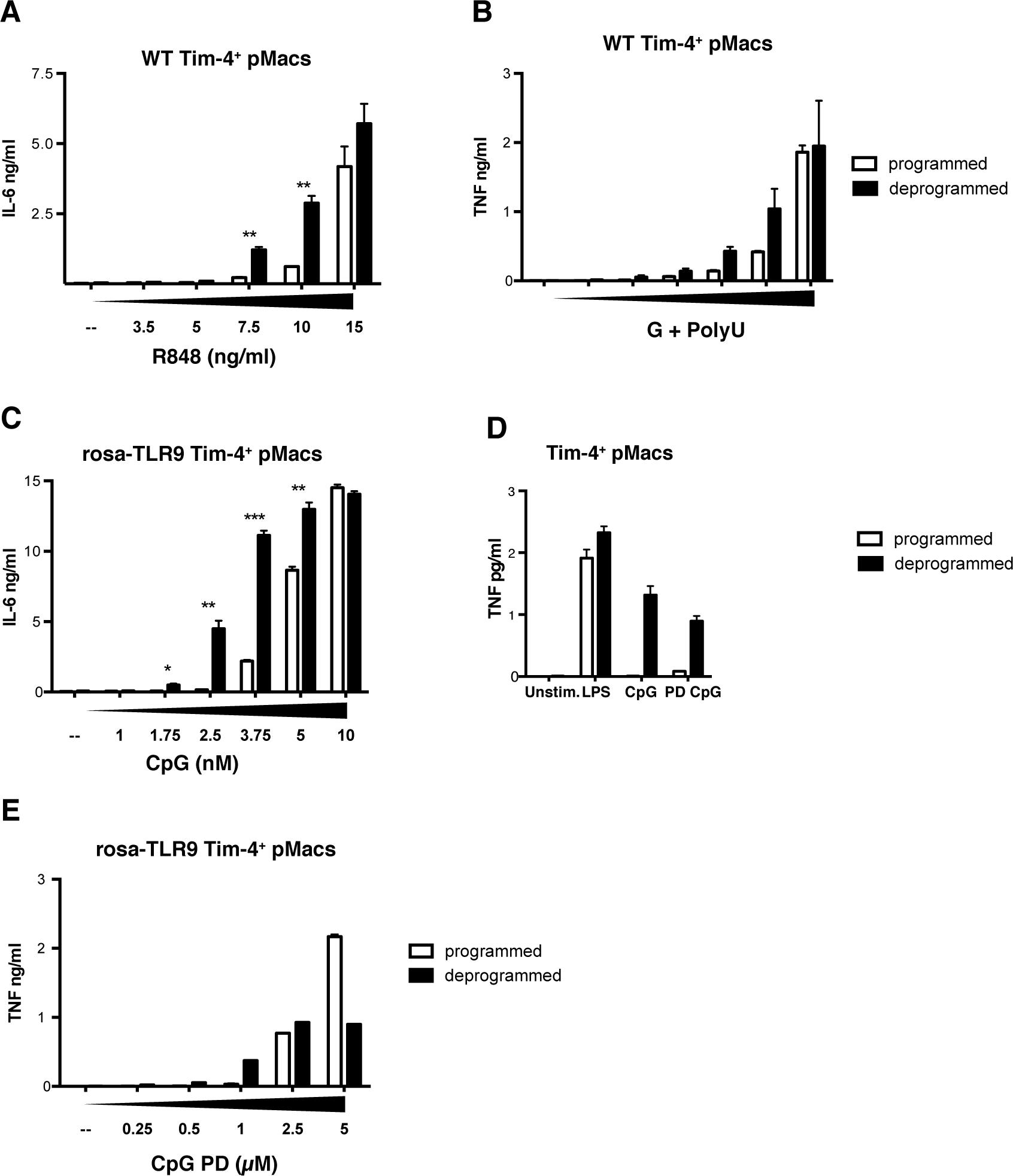
Programmed macrophages have a higher activation threshold for TLR7 and TLR9 responses, related to Figure 4 and Figure 5. **(A)** Deprogrammed Tim-4^+^ pMacs respond to lower doses of TLR7 ligand. Programmed and deprogrammed WT Tim-4^+^ pMacs were stimulated with increasing doses of R848. IL-6 was measured by CBA. Data are representative of at least three independent experiments. **(B)** Deprogrammed Tim-4^+^ pMacs from rosa-TLR9 mice respond to lower doses of TLR9 ligand. Programmed and deprogrammed Tim-4^+^ pMacs from rosa-TLR9 mice were stimulated with increasing doses of CpG ODN. IL-6 was measured by CBA. Data are representative of at least three independent experiments. **(C)** Deprogrammed Tim-4^+^ pMacs respond to lower doses of the TLR7 ligand G + PolyU. Programmed and deprogrammed WT Tim-4^+^ pMacs were stimulated with 2 fold dilutions of G + PolyU with a starting concentration of 0.1 mM G + 10μg/ml PolyU. TNF was measured by CBA. Data are representative of two independent experiments. **(D)** Deprogrammed Tim-4^+^ pMacs from rosa-TLR9 mice respond to lower doses of the TLR9 ligand phosphodiester backbone CpG. Programmed and deprogrammed Tim-4^+^ pMacs from rosa-TLR9 mice were stimulated with increasing doses of PD CpG ODN. TNF was measured by CBA. Data are representative of two independent experiments.

**Figure S6.**
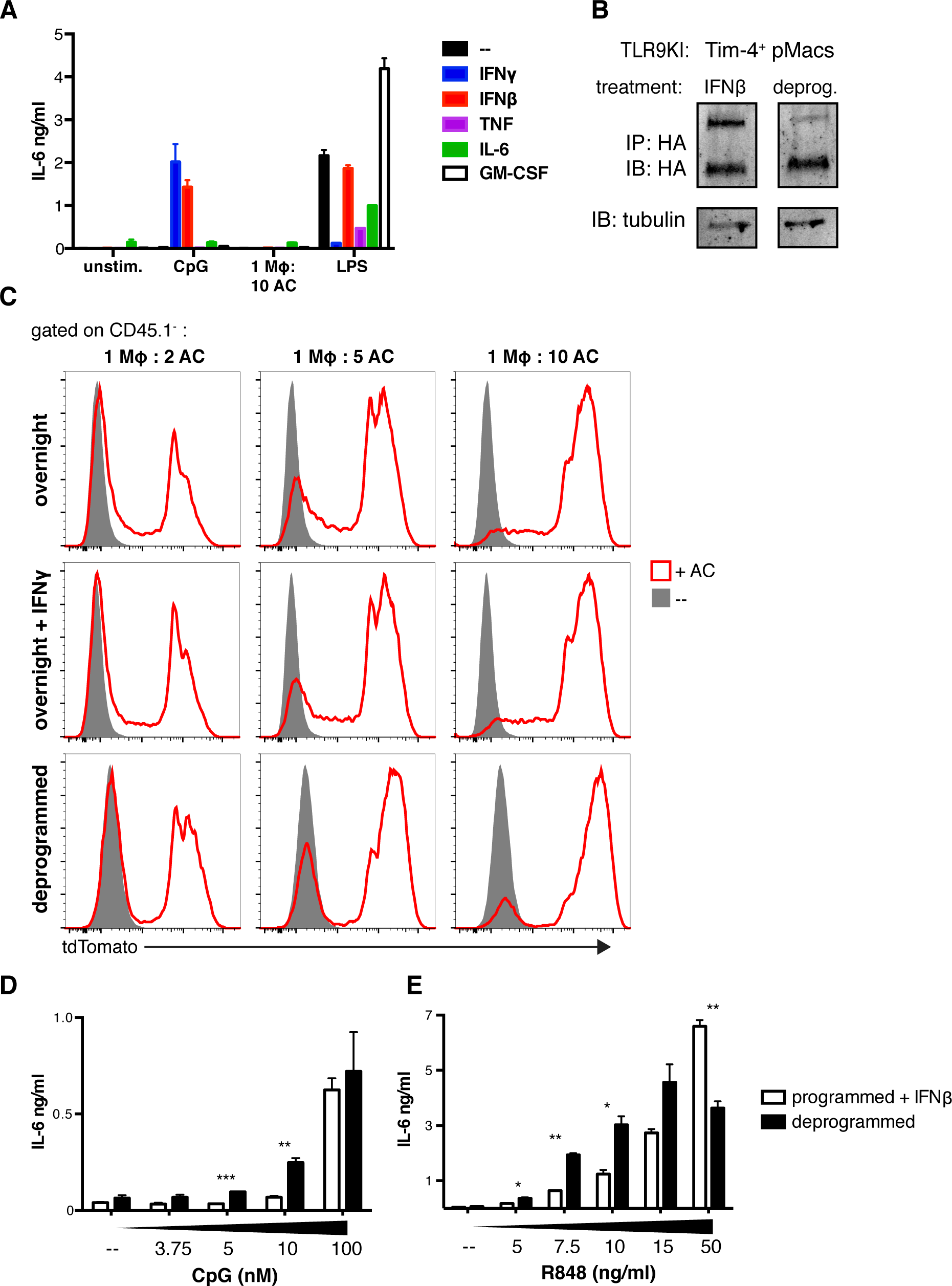
IFNγ treated macrophages are adept at AC engulfment, related to Figure 6. **(A)** IFNγ and IFNβ induce responses to TLR9 ligands in Tim-4^+^ pMacs. Isolated Tim-4^+^ pMacs were treated overnight with the indicated cytokines before stimulation with TLR ligands or ACs. Concentrations of cytokines used were 10ng/ml IFNγ, 20U/ml IFNβ, 5ng/ml TNF, 50ng/ml IL-6, or 5ng/ml GM-CSF. Cytokine responses were measured by CBA. Data are representative of at least three independent experiments. **(B)** IFNβ treated and deprogrammed Tim-4^+^ pMacs express similar levels of TLR9. Tim-4^+^ pMacs from TLR9KI mice were cultured overnight ± IFNβ or for 60 hours, and TLR9 levels in lysates were measured by anti-HA immunoprecipitation and immunoblot. An anti-tubulin immunoblot was performed on each lysate as a reference. Shown are the indicated lanes of the same membrane. Data are representative of two independent experiments. **(C)** Tim-4^+^ pMacs were isolated from WT mice and cultured overnight ± IFNγ or for 60 hours then incubated with CD45.1^+^tdTomato^+^ ACs at indicated ratios and analyzed by flow cytometry. **(D)** IFNβ treated Tim-4^+^ pMacs maintain a high threshold for TLR9 responses. Isolated programmed IFNβ treated and deprogrammed untreated Tim-4^+^ pMacs were stimulated with increasing doses of CpG. IL-6 was measured by CBA. Data are representative of two independent experiments. **(E)** IFNβ treated Tim-4^+^ pMacs maintain a high threshold for TLR7 responses. Isolated programmed IFNβ treated and deprogrammed untreated Tim-4^+^ pMacs were stimulated with increasing doses of R848. IL-6 was measured by CBA. Data are representative of two independent experiments.

**Figure S7.**
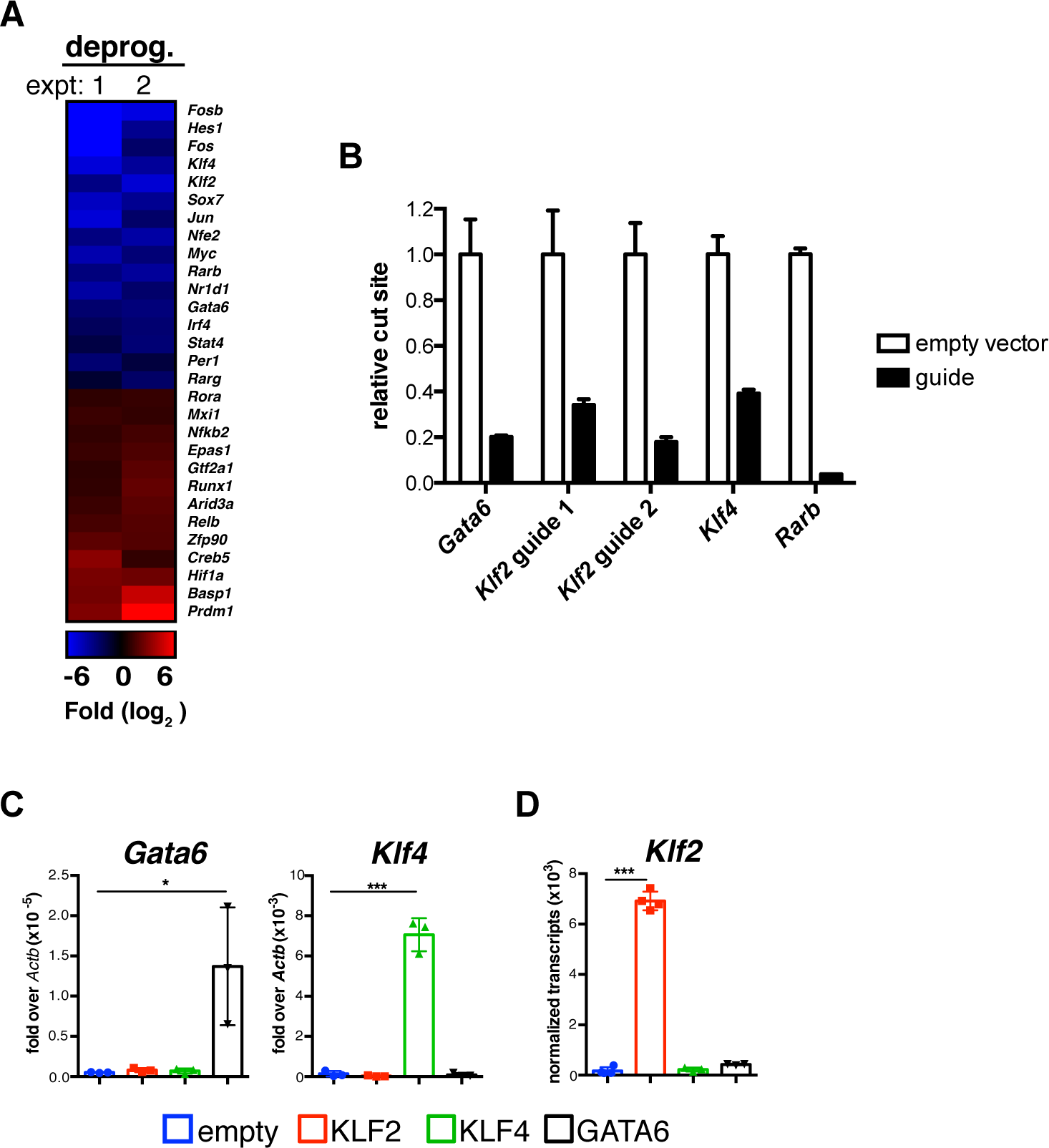
KLF2 and KLF4 imprint an AC-clearance program on macrophages, related to Figure 7. **(A)** Changes in transcription factor expression associated with deprogramming. RNA was isolated from programmed and deprogrammed WT Tim-4^+^ pMacs and analyzed by RNA sequencing. The heat map indicates transcription factors that are upregulated or downregulated at least 2-fold (p < 0.05) in deprogrammed Tim-4^+^ pMacs relative to programmed Tim-4^+^ pMacs. Results of two independent experiments are shown. Replicates have been averaged so only one value per experiment is shown **(B)** qPCR was performed using primers that bind cas9 cut site to determine efficiency of knock out in BMMs from cas9 overexpressing mice that were transduced with vectors encoding guide RNAs targeting the indicated transcription factors. Genomic DNA from BMMs was isolated and qPCR was performed. Data was normalized to an unrelated site in genomic DNA. **(C and D)** BMMs were transduced with retrovirus encoding KLF2, KLF4, GATA6 or empty vector. (C) The expression of GATA6 and KLF4 was determined by qPCR and is expressed relative to *Actb* mRNA. (D) The expression of KLF2 were quantified using a Nanostring nCounter. Data are the combined results of three independent experiments. n = 3-4 per group from independent experiments.

## Experimental procedures

### Mice

Mice were housed under specific pathogen-free conditions at the University of California, Berkeley. All mouse experiments were performed in accordance with the guidelines of the Animal Care and Use Committee at UC Berkeley. Unless noted mice were analyzed at 6-12 weeks of age. C57BL/6, C57BL/6 CD45.1^+^ (stock #002014), rosa26 ^stop-flox-tdTomato^ (stock #007909), LysM-cre (stock #004781), and EIIa-cre (stock #003724) mice were obtained from Jackson laboratories. Unc93B1^−/-^ mice were generated by the trans-NIH Knock-Out Mouse Project (KOMP) and obtained from the KOMP Repository (www.komp.org). Tim-4^−/-^ mice were obtained from Genentech (Wong et al., 2010). LysMcre^+/-^KLF2^fl/fl^ mice were obtained from Jerry Lingrel (University of Cincinnati) (Weinreich et al., 2009).

TLR7 reporter mice were generated by constructing a targeting vector encoding a FLAG tag on the 3’ end of the TLR7 gene followed by IRES-tdTomato sequence. This vector was electroporated into C57BL/6-derived embryonic stem cells by the Mouse Biology Program at UC Davis. The vector also introduced a loxP-flanked neomycin resistance cassette. Targeting was assessed by southern blot and correctly targeted ES cells were injected into ICR/CD1 blastocysts. Chimeric males were mated with C57BL/6 background EIIA-cre females to remove the neomycin resistance cassette. The TLR9 reporter mice were generated in a similar manner except the construct encoded an HA tag on the 3’ end of the TLR9 gene followed by an IRES-EGFP sequence. Rosa^stop-flox-TLR9^ mice will be described in detail elsewhere. Briefly, a floxed region (containing eGFP, a Neo cassette, and a transcriptional stop sequence) followed by an HA tagged TLR9 was inserted into the Rosa26 locus. The construct was generated by cloning cDNA encoding TLR9 with a C-terminal HA tag into the pBigT vector to introduce a loxP-flanked cassette with a transcriptional stop sequence. The resulting construct was cloned into the ROSA26 eGFP DTA plasmid, thereby replacing DTA with HA tagged TLR9. pBigT invloxP (pBigT) and ROSA26 eGFP DTA were a kind gift from Martinez-Barbera (Ivanova et al., 2005). The linearized targeting construct was electroporated into C57BL/6 ES cells. Correctly targeted ES clones were identified by southern blot and correctly targeted clones were injected into blastocysts. Chimeric mice were mated to achieve germline transmission.

Chimeric mice were generated by irradiation of C57BL/6 with 1000rads. Mice were reconstituted with 1x10^7^ bone marrow cells (80% C57BL/6 and 20% CD45.1 + tdTomato^+^). Mice were analyzed 10-12 weeks after cell transfer.

### Tissue harvest

Cells from the peritoneal and pleural cavities were recovered by lavage with ice cold PBS. Spleens and lymph nodes were digested with collagenase XI (Sigma #C9697) with DNAse I (Sigma #D4513) for 30min and single cell suspensions were generated by mechanical disruption. Perfused lungs were digested with collagenase XI with DNAse I for 45min, single cell suspensions were generated by mechanical disruption through a 100um filter, cells were resuspended in 44% isotonic percoll (GE healthcare #17-0891- 01), underlayed with 67% percoll, and spun at 1550xg without brake. Cells from the interface were collected for analysis. Perfused livers were digested with collagenase VIII (Sigma #C2139) with DNase I for 45min, cells were resuspended in Hank’s buffered salt solution and centrifuged at 30xg for 3min. After low speed centrifugation cells in suspension were collected and underlayed with a solution of 25% isotonic Percoll and 50% Percoll, then spun for 15min at 1800xg without brake. Cell counts were obtained using Count Bright absolute counting beads (Life technologies #C36950).

### Isolation of peritoneal Macrophages

Peritoneal cells were recovered by lavage with 5ml of ice cold PBS. For RNAseq and western blot experiments B cells were first depleted using anti-CD19-biotin antibody and biotin binder dynabeads (ThermoFisher #11047). Tim-4^+^ cells were isolated using anti-TIM-4 antibody (clone RMT4-54, BioLegend) and anti-rat IgG microbeads (Miltenyi #130-048-501). F4/80^+^ cells were isolated using anti-F4/80 antibody (clone BM8, eBioscience and anti-rat IgG microbeads.

### Cell culture

HEK293 cells were cultured in DMEM supplemented with 10% (v/v) fetal calf serum, L-glutamine, penicillin-streptomycin, sodium pyruvate, and HEPES pH 7.2 (all supplements and media were purchased from Gibco). BMMs were differentiated for seven days in RPMI complete media (RPMI-1640 supplemented with 10% (vol/vol) fetal calf serum, L-glutamine, penicillin-streptomycin, sodium pyruvate, and HEPES pH 7.2) supplemented with M-CSF containing supernatant from 3T3-CSF cells.

Unless noted peritoneal macrophages *ex vivo* for 60 hours, as well as overnight controls, were cultured in 25% (v/v) omentum supernatant in RPMI complete media. Omentum supernatant was generated by harvesting omenta from C57BL/6 mice and culturing the omenta in RPMI complete media at 1ml/omenta overnight. Supernatant was collected, filtered through a 0.22um filter, and frozen at -80**°**C for future use. Lung and pleural cells were harvested as described above and plated on non-tissue culture treated plates in RPMI complete media.

For overnight cytokine treatment, cytokines were added to RPMI complete media containing 25% omentum culture supernatant and incubated with macrophages for 16 hours. Concentrations of cytokines used were 10ng/ml IFNγ, 20U/ml IFNβ, 5ng/ml TNF, 50ng/ml IL-6, or 5ng/ml GM-CSF. For dose curve comparison and corresponding western blot IFNβ was used at 100U/ml. (IFNγ – Tonbo #21-8311-U020, IFNβ – Pbl interferon source #12401-1, TNF – aa84-235 R&D #410-TRNC, IL-6 R&D #406-ML-025, GM-CSF Tonbo #21-8331-U020).

### Apoptotic cell generation and engulfment

Thymi were harvest from WT or CD45.1^+^ rosa-tdTomato mice and single cell suspensions were generated by mechanical disruption through a 70um filter. Cells were irradiated with 600rad and incubated in RPMI complete media at 37°C for 4 hours. Apoptotic cells were then incubated with macrophages at the indicated ratios. For experiments examining AC engulfment capabilities macrophages were allowed to engulf ACs for 60min. For AC injection experiments 0.5e6 CD45.1^+^ rosa-tdTomato ACs were injected intraperitoneally per mouse; one hour after injection peritoneal cells were harvested as above.

### Stimulations

Cells were plated in RPMI complete media. Tissue macrophages were stimulated directly *ex vivo*, after overnight culture, or after 60 hours in culture. Cells were stimulated with TLR ligands (LPS, CpG-B ODN 1668, R848 all from Invivogen), or ACs. For analysis of secreted cytokines, supernatant was collected 7 hours after stimulation. For intracellular cytokine staining, 30 minutes after stimulation brefeldin A (GolgiPlug, BD Biosciences) was added to cells before incubation for another 4 hours.

### Flow cytometry and Antibodies

Dead cells were excluded using a fixable live/dead stain (Life technologies) or DAPI (Life Technologies) and all stains were carried out in PBS containing 2% FBS (v/v) and 2mM EDTA including anti-CD16/32 Fc blocking antibody (2.4G2, UCSF monoclonal antibody core) and normal mouse serum (Sigma). Cells were stained for 20min at 4**°**C with antibodies (see table 1).

**Table 1:** Antibodies.

| **Antibody** | **Clone** | **Vendor** |
| --- | --- | --- |
| B220 | RA3-6B2 | eBioscience |
| CD11c | N418 | BioLegend |
| CD19 | 6D5 | BioLegend |
| CD3ε | 145-2C11 | Tonbo |
| CD11b | M1/70 | BioLegend |
| CD45.1 | A20 | BioLegend |
| CD45.2 | 104 | BioLegend |
| CD64 | X54-5/7.2 | BioLegend |
| F4/80 | BM8 | Tonbo |
| F4/80 | Cl:A3-1 | Biorad |
| Ly6C | HK1.4 | BioLegend |
| Ly6G | 1A8 | BioLegend |
| MHC II | M5/114.15.2 | eBioscience |
| Siglec-F | E50-2440 | BioLegend |
| Tim-4 | RMT4-54 | BioLegend |
| TNF | MP6-XT22 | BioLegend |
| GATA6 | D61E4 | Cell Signaling |
| pERK1/2 biotin | D13.14.4E | Cell Signaling |
| HA antibody | 3F10 | Roche |
| tubulin | DM1A | Calbiochem |

**Table 2:**
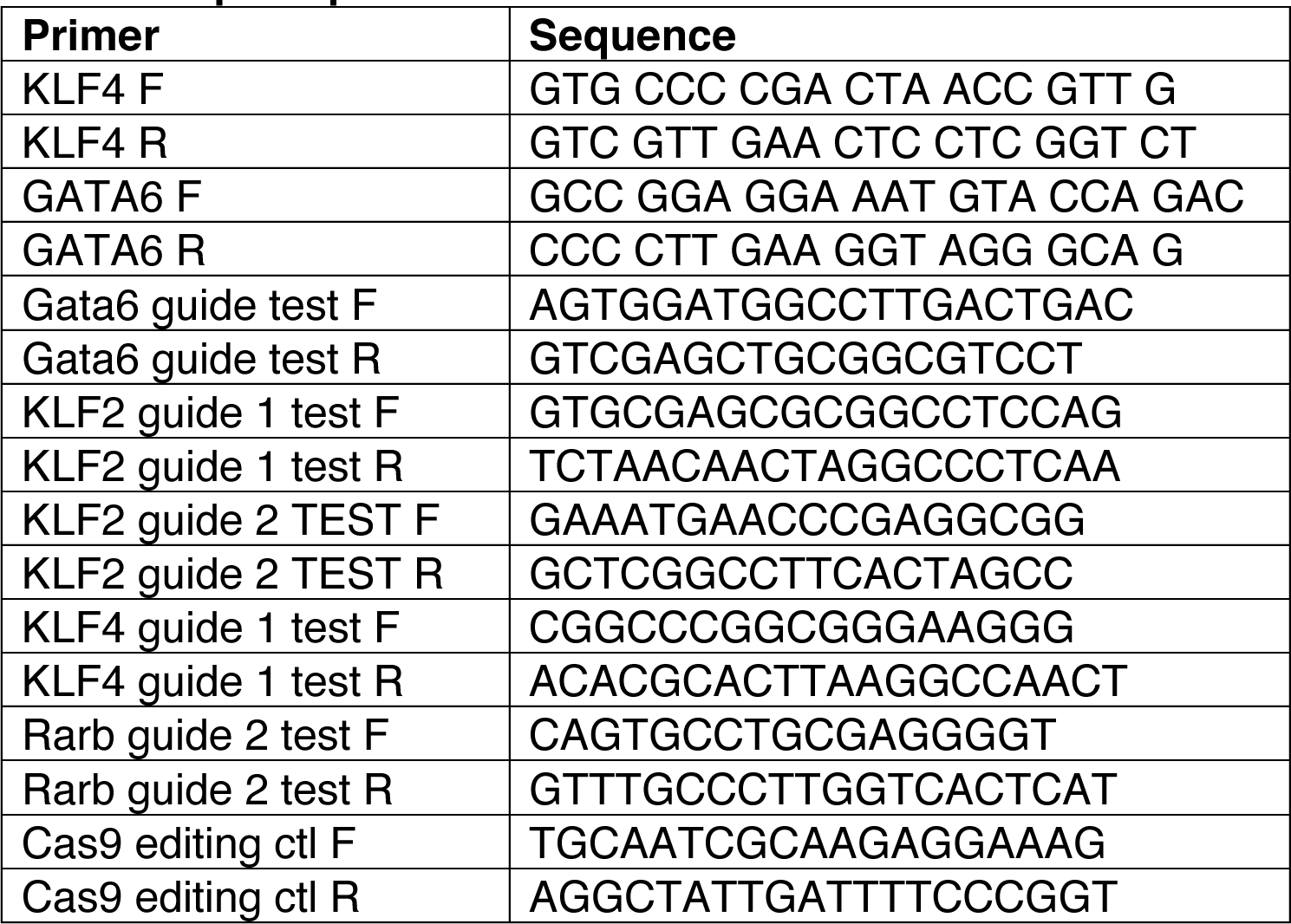
qPCR primers.

**Table 3:**
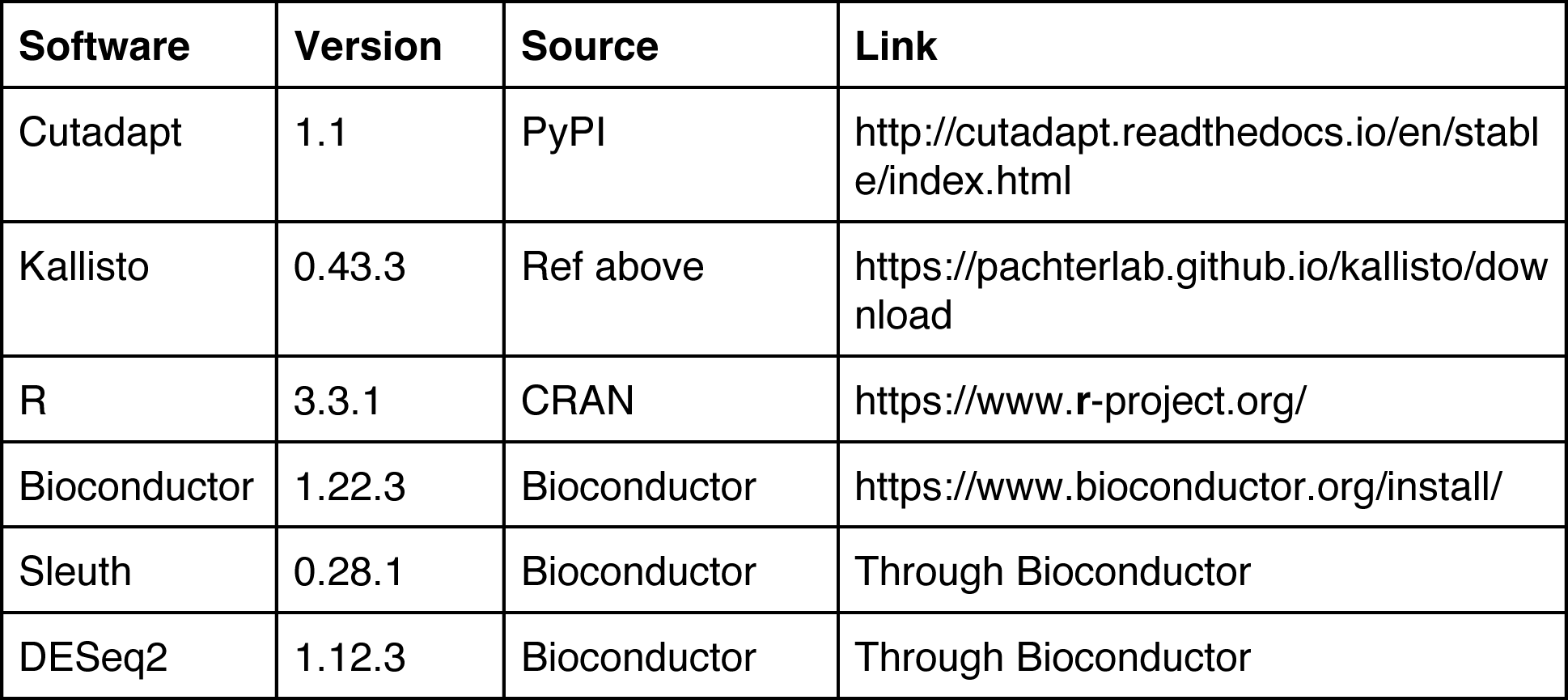
Software used for RNA sequencing analysis.

| **Software** | **Version** | **Source** | **Link** |
| --- | --- | --- | --- |
| Cutadapt | 1.1 | PyPI | http://cutadapt.readthedocs.io/en/stable/index.html |
| Kallisto | 0.43.3 | Ref above | https://pachterlab.github.io/kallisto/download |
| R | 3.3.1 | CRAN | https://www.r-project.org/ |
| Bioconductor | 1.22.3 | Bioconductor | https://www.bioconductor.org/install/ |
| Sleuth | 0.28.1 | Bioconductor | Through Bioconductor |
| DESeq2 | 1.12.3 | Bioconductor | Through Bioconductor |

For intracellular TNF or GATA6 staining cells were permeabilized with Fix/Perm buffer (BD or eBioscience respectively) for 20min at 4**°**C. Cells were then stained with antibodies. For pERK staining cells were immediately fixed after stimulation with 1.6% PFA at room temperature, then washed and resuspended in ice-cold methanol. After overnight incubation at -20°C, cells were stained with p-ERK1/2 biotin followed by Steptavidin-BV421 (BioLegend). All cells were analyzed on an LSRII or LSR Fortessa (BD Biosciences), and data was analyzed with FlowJo (TreeStar).

Activated Caspase-3/7 was measured using CellEvent Caspase-3/7 Green Flow Cytometry Assay (ThermoFisher), according to the manufacturer’s instructions.

### BMM transfers

For peritoneal transfer experiments 1e6 differentiated CD45.1^+^ BMMs were injected into the peritoneum of CD45.2^+^ mice. Peritoneal cells were harvested 2 or 3 weeks after cell transfer and stimulated with TLR ligands as described above. For lung transfer experiments differentiated 2e6 CD45.1^+^ Tomato^+^ BMMs were intranasally administered on three consecutive days. Lungs were harvested, as described above, 5.5 weeks after cell transfer. For CFSE proliferation experiments CD45.1^+^BMMs were labeled with 5µM CFSE in PBS for 10min at 37°C prior to transfer.

### Transcription factor overexpression

For retroviral transduction of BMMs a mouse stem cell virus (MSCV)-based retroviral vector was used that expressed the gene of interested followed by IRES-puromycin resistance gene-T2A-EGFP. VSV-G-pseudotyped retrovirus was made in GP2-293 packaging cells. GP2-293 cells were transfected with retroviral vectors and pVSV-G using Lipofectamine LTX reagent (ThermoFisher #15338100). 24 hr post-transfection, cells were incubated at 32°C. 48 hr post-transfection viral supernatant was used to infect bone marrow cells in RPMI complete media containing M-CSF. Bone marrow cells and viral supernatant were spun at 1250xg for 90 min at 32°C then incubated overnight at 32°C. Bone marrow cells were re-transduced with fresh viral supernatant the following day. Two days after last transduction puromycin selection was added for three days.

For lentiviral overexpression a vector with a pHAGE backbone was used that expressed GATA6 followed by IRES-mCherry. Lentiviral particles were produced in 293T cells by transfecting with the lentiviral plasmid and the helper constructs psPAX2 and pCMV-VSV-G. Lentivirus was concentrated by filtering cell supernatant with 0.45µm filter then centrifuging at 47,850xg. Viral pellets were resupended in PBS. 0.5e6 transduction units were injected per mouse. One week after injection peritoneal cells were harvested.

### Cas9 genome editing of BMMs

The lentiviral gRNA plasmid pKLV-U6gRNA(BbsI)-PGKpuro2ABFP (Addgene #50946) was used. The sequences of the guide RNA target sites are as follows with the protospacer adjacent motif underlined: GATA6 - ACCGCCTCGGCGTCGAGCTG<u>CGG</u>, KLF2 guide 1 – TTCGCCAGCCCGTGCGAGCG<u>CGG</u>, KLF2 guide 2 - CTGGCCGCGAAATGAACCCG<u>AGG</u>, KLF4 – CTCCACGTTCGCGTCCGGCC<u>CGG</u>, RARβ - TATGGCGTCAGTGCCTGCGA<u>GGG</u>. BMMs from rosa-Cas9 mice (Jackson Laboratory) were transduced with lentivirus and selected with puromycin.

### Pristane

Mice were intraperitoneally injected with 0.5ml of pristane (Sigma #P2870) that had been filtered through a 0.22um filter. Mice were analyzed after two weeks or after nine months.

### Western Blot Analysis

Cells were lysed in RIPA buffer (50mM Tris pH 7.4, 150mM NaCl, 1mM EDTA, 0.5mM EGTA, 1% NP-40, 1% DOC, 0.1% SDS) containing protease inhibitors (Roche #05 892 791 001). Cell lysates were immunoprecipitated with anti-HA matrix (Roche #11573000). After washing in RIPA buffer, matrix beads were boiled in SDS-PAGE loading buffer. Protein was run on a 4-15% gel (Biorad #4561083) and transferred to Immobilon-FL membrane (Millipore #IPFL00010). After blocking, membranes were probed with anti-HA antibody, followed by anti-rat-680 secondary (Life Technologies #A21096) or probed with anti-tubulin, followed by anti-mouse-800 secondary (Li-Cor #926-32210). Images were scanned using a Licor Odyssey. Images were quantified using ImageJ.

### Enzyme-linked immunosorbent assay (ELISA) and CBA assay

Cytokines were measured using a Cytometric Bead Array assay according to the manufacturer’s instructions (BD Biosciences).

For anti-dsDNA ELISAs NUNC Maxisorp plates (eBioscience) were coated first with poly-l-lysine (Sigma #P-6516) at 50µg/ml overnight at -20°C, washed, then coated with 100µg/ml dsDNA (Calf thymus DNA Sigma #D45220) overnight at 4°C. Plates were then blocked with PBS + 1% BSA (w/v) + 5% goat serum (v/v) at room temperature for four hours before serum samples diluted 1:500 in PBS + 1% BSA (w/v) + 5% goat serum (v/v) were added and incubated overnight at 4°C. Secondary anti-IgG all-biotin (Jackson #115-065-205) was used at 1:10000 followed by Streptavidin-HRP (BD Biosciences #554066). Plates were developed with 1mg/ml OPD (Sigma #P6912) in Citrate Buffer (PBS with 0.05M NaH2 PO4 and 0.02M Citric acid) with HCl acid stop.

### RNA sequencing

Total RNA was prepared from directly *ex vivo* or deprogrammed (60 hour cultured) Tim-4^+^ pMacs using RNAzol (Molecular Research Center #RN 190), DNAse treated (Turbo DNase from Ambion #AM2238) and purified using RNA clean and concentrator columns (Zymo #R1015). 1 ug of total RNA was subsequently used for rRNA depletion using the RiboZero depletion kit to prepare RNA-Seq library by using the TruSeq RNA sample prep kit (Illumina) according to manufacturer’s instructions. 50- cycle single-end RNA sequencing was performed on a HiSeq 4000 (Illumina). Sequenced reads were quality-trimmed (cut-off of 30) with cutadapt and aligned to the mouse transcriptome (GRCm38, release 79) with Kallisto (Bray et al., 2016). Using R and Bioconductor packages (see list in table), counts for each transcript were extracted using Sleuth, collapsed at the gene level, and analyzed for differential expression using DESeq2. Q-values (Benjamini-Hochberg correction) lower than 0.05 were considered significant. Heat maps were generated using MeV (Multiple Experiment Viewer) software.

For the AC stimulation experiment AC derived RNA was analyzed as well as RNA from macrophages ± ACs. Peritoneal macrophages were stimulated with ACs immediately *ex vivo* (programmed) or after 60 hours in culture (deprogrammed). Two hours after addition of ACs RNA was prepared as above. Genes whose expression was greater than two fold higher in RNA isolated from ACs alone relative to macrophage RNA were considered AC-derived and excluded.

### Quantitative PCR

Cells were lysed in RNAzol (MRC) and RNA was purified according to manufacturer’s instructions and concentrated using a column (Zymo #R1015). cDNA was prepared with iScript cDNA synthesis kit (Biorad), and quantitative PCR was performed with SYBR Green (Biorad) on a StepOnePlus thermocycler (Applied Biosystems). Primer sequences for GATA6 and KLF4 were obtained from PrimerBank and synthesized by IDT.

For analysis of cas9 genome editing genomic DNA from BMMs was prepared using a genomic DNA isolation kit (Biorad #732-6340). Primers were designed to bind the cut site and normalized using control primers that bind an unrelated site (see table).

### mRNA transcript count using Ncounter

The expression levels of a set of genes associated with tissue macrophages, AC clearance, or TLR signaling were analyzed using the NanoString nCounter Analysis System (NanoString Technologies). Raw counts of samples were normalized according to the manufacturer’s recommendations using reference genes as internal controls (Cltc, Gapdh, Gusb, Hprt, Pg1, and Tubb5). Normalization was performed using nSolver Analysis Software v3.0 (NanoString Technologies)

### Statistical Analysis

Statistical analysis was performed with the Prism software (GraphPad software). P-values were determined using unpaired two-tailed Students’s *t*-test. Where noted *t*- tests were performed on log transformed data to account for the non-normal distribution of the data. *p < 0.05, **p < 0.01, ***p < 0.001.

